# Sylites: Multipurpose markers for the visualization of inhibitory synapses

**DOI:** 10.1101/2021.05.20.444651

**Authors:** Vladimir Khayenko, Clemens Schulte, Sara L. Reis, Orly Avraham, Cataldo Schietroma, Rafael Worschech, Noah F. Nordblom, Sonja Kachler, Carmen Villmann, Katrin G. Heinze, Andreas Schlosser, Ora Schueler-Furman, Philip Tovote, Christian G. Specht, Hans Michael Maric

## Abstract

We introduce Sylites – small and versatile fluorogenic affinity probes for high-contrast visualization of inhibitory synapses. Having stoichiometric labeling and exceptional selectivity for neuronal gephyrin, a hallmark protein of the inhibitory post-synapse, Sylites enable superior synapse staining compared with antibodies. Combined with super-resolution microscopy, Sylites allow precise nanoscopic measurements of the synapse. In brain tissue, Sylites reveal the three-dimensional distribution of inhibitory synapses within just an hour.

## Main

Reliable markers that visualize synapses, and, by extension, neural circuits, have great value for clinical and fundamental neuroscience^1^. An integral component of inhibitory synapses is gephyrin, a highly abundant scaffold protein that stabilizes glycine and GABA_A_ receptors^2,3^. Gephyrin serves as a universal marker of the inhibitory synapse and its concentration at the post synaptic density closely correlates with the number of inhibitory receptors and the synaptic strength^4–6^. Gephyrin is commonly visualized using antibodies^7^ or recombinant techniques that tag gephyrin with fluorescent proteins^8^, but these approaches come with caveats: recombinant proteins are prone to overexpression, their use in complex organisms is challenging and they cannot be applied if wild-type species are to be studied. Antibodies, on the other hand, do not require genetic manipulation and are easily applicable in fixed samples; however, their large size and the tendency to crosslink affect the labeling performance in complex samples^9^. Here we introduce selective high affinity stoichiometric probes for gephyrin that enable high-contrast visualization of synapses in cell cultures and tissue and deliver accurate super-resolution measurements.

The first probe that exploited a definite feature of synaptic gephyrin, a universal receptor binding pocket in the gephyrin E domain, was TMR2i^10^. This probe was derived from the intracellular loop of the glycine receptor (GlyR) β subunit, a natural ligand of this docking site, as are multiple members of the synaptic GABA_A_R subtypes^11,12^. TMR2i did bind gephyrin but gave low-contrast labeling and was not suitable for nanoscopy, such as direct stochastic optical reconstruction microscopy (dSTORM).

We systematically optimized both the probe architecture and the gephyrin binding sequence to target the native protein (Fig.S1, Tables S2,4,5), and synthesized an array of fluorescent probes. We then performed an imaging-based evaluation of the probes for the binding of gephyrin in fixed cells (Fig.S2, Fig.S3). SyliteM, a monomeric probe with strictly linear gephyrin labeling (Fig.S2), and SyliteD, a dimeric probe (Fig.1a) with higher affinity and contrast, displayed the highest target to off-target labeling ratios and a strong linear relationship with gephyrin (Fig.S2).

**Figure 1.**
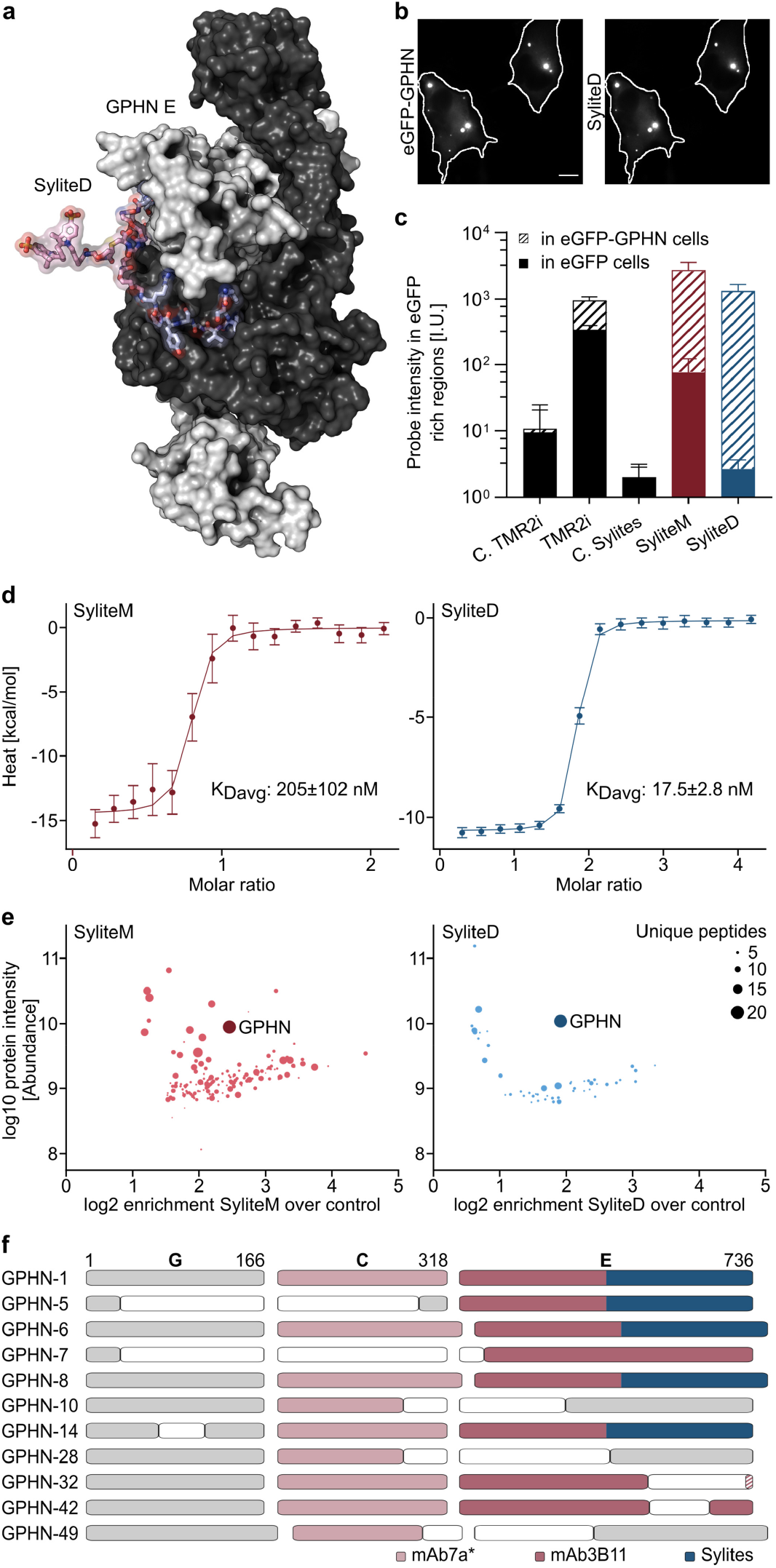
Sylites selectively and stoichiometrically bind gephyrin, enabling high contrast imaging. **a.** Rosetta FlexPepDock structural model of SyliteD bound to gephyrin (GPHN) E domain dimer. In blue is the binding sequence, in pink is the linker and the fluorophore, the two gephyrin E domains in black or white. **b.** Left: Fixed COS-7 cells expressing eGFP-gephyrin, Right: SyliteD 50 nM staining of the fixed sample. Scale bar 10 μm. **c.** Labeling contrast of the peptide-based probes. The Y axis represents log average signal intensity of the probe signal from eGFP-rich regions of the COS-7 cells. SyliteM and SyliteD have ~35 and ~500 target to off-target labeling ratio, respectively, TMR2i has a target to off-target labeling ratio of ~3. C.TMR2i represents red spectrum readouts of unlabeled cells. C.Sylites represents far-red spectrum readouts of unlabeled cells. n≥8. Mean±SD. **d.** ITC measured heat signature of SyliteM and SyliteD titrated with gephyrin E domain. Both probes exhibit nanomolar affinity, with SyliteD having 10-fold affinity increase over the monomer. **e.** Sylites selectively pull gephyrin. Quantitative mass spectrometric analysis of SyliteM/D pull-downs. Non-fluorescent versions of SyliteM and SyliteD were used to pull down proteins from mouse brain homogenate and the protein fractions were subsequently digested and analyzed with LC-MS/MS. The size of the circle corresponds to the number of unique peptides identified for the specific protein. Left: SyliteM pulls down additional proteins that have high intensity and are abundant in its pool, although gephyrin is the most prominent. Right: gephyrin is a single protein with high abundance, selectivity and strong representation in the SyliteD pull-down. **f.** Sylites bind synaptic gephyrin. Probe interaction with gephyrin isoforms. The isoform GPHN-1 represents the primary structure of gephyrin^19^. The protein consists of 3 regions: G domain, C unstructured linker region, E domain, where the receptor-binding pocket is located. Blank boxes indicate deletions, elongated boxes – additions, striped boxes – substitutions. The peptide probes (blue) bind to isoforms that can form a receptor binding pocket of the E domain. The antibodies bind both competent and incompetent receptor clustering isoforms. mAb7a (rose) binds a short linear *Ser270 phosphorylated epitope in the “C” linker region, while mAb3B11 (raspberry) interacts with an epitope in the E domain.

Sylites showed a near-complete correlation with gephyrin in COS-7 cells expressing recombinant eGFP-gephyrin, and no correlation with cells expressing soluble eGFP. Notably, the target to off-target labeling ratios of SyliteM and SyliteD were ~35 and ~500, respectively, approximately 10 and 150-fold higher than those of TMR2i (Fig.1b,c, Fig.S3). Using isothermal titration calorimetry (ITC) with gephyrin E domain, we determined a K_d_ of 17.5±2.8 nM for the dimeric SyliteD and a K_d_ of 205±102 nM for the monomeric SyliteM, indicating high probe affinity, and confirming the stoichiometric binding of 1:1 for SyliteM and 1:2 for SyliteD (Fig.1d), in line with their monomeric and dimeric design. Lastly, mass-spectrometric determination of the interactomes of Sylites confirmed their target selectivity. Gephyrin was the only protein with high abundance, high selectivity and multiple unique peptide fragments binding to the dimeric probe. The monovalent probe retained some additional proteins other than gephyrin, consistent with its somewhat lower target to off-target labeling ratio.

Gephyrin is a multifunctional protein with numerous isoforms and post-translational modifications^13^. Comparison of the binding profiles to eleven major gephyrin isoforms expressed in HEK293 cells (Fig.1f, Fig.S4) reveals that Sylites, but not the tested antibodies, exclusively label gephyrin isoforms that have GlyR and GABAA receptor binding capacity. This indicates that Sylites are ideally suited to detect synapses and to quantify functionally relevant receptor binding sites. Interestingly, no gephyrin labeling was observed with the widely used mAb7a antibody in HEK293 cells. Microarray profiling of mAb7a binding (Fig.S5) confirms that in contrast to Sylites, mAb7a depends on a phosphorylated (pSer270) epitope in the linker region of gephyrin^14^. Thus, mAb7a labels only a sub-population of synaptic gephyrin isoforms and phosphorylation variants.

We next used the probes to study the structure and distribution of inhibitory synapses in brain sections and cultured neurons using conventional and super-resolution microscopy. In cortical neurons expressing gephyrin-mEos2 fluorescent protein chimera, we observed high contrast visualization of densely packed gephyrin clusters at synapses and a near complete correlation of Sylite staining with gephyrin (Fig.2a,b). Target specificity was confirmed by Sylite overlap with endogenous antibody-labeled gephyrin in wild-type cortical and hippocampal neurons (Fig.S6). Linear regression analysis of fluorescent intensities of synaptic mEos2-gephyrin puncta with correspondent Sylite or mAb7a-stained puncta revealed a 3- and 2-time narrower prediction interval for SyliteM and SyliteD, respectively, compared with mAb7a, indicating a much closer relation between Sylite and mEos2-gephyrin signals (Fig.2c). The higher scattering observed with mAb7a, suggests that the antibody staining exhibits non-linear scaling with synaptic gephyrin, in agreement with our previous findings on the selectivity of mAb7a for a specific, phosphorylated variant of gephyrin (Fig.S5). Taken together, our data demonstrate a linear, stoichiometric relationship between Sylites and gephyrin, making them suitable for quantitative microscopy^15^.

**Figure 2.**
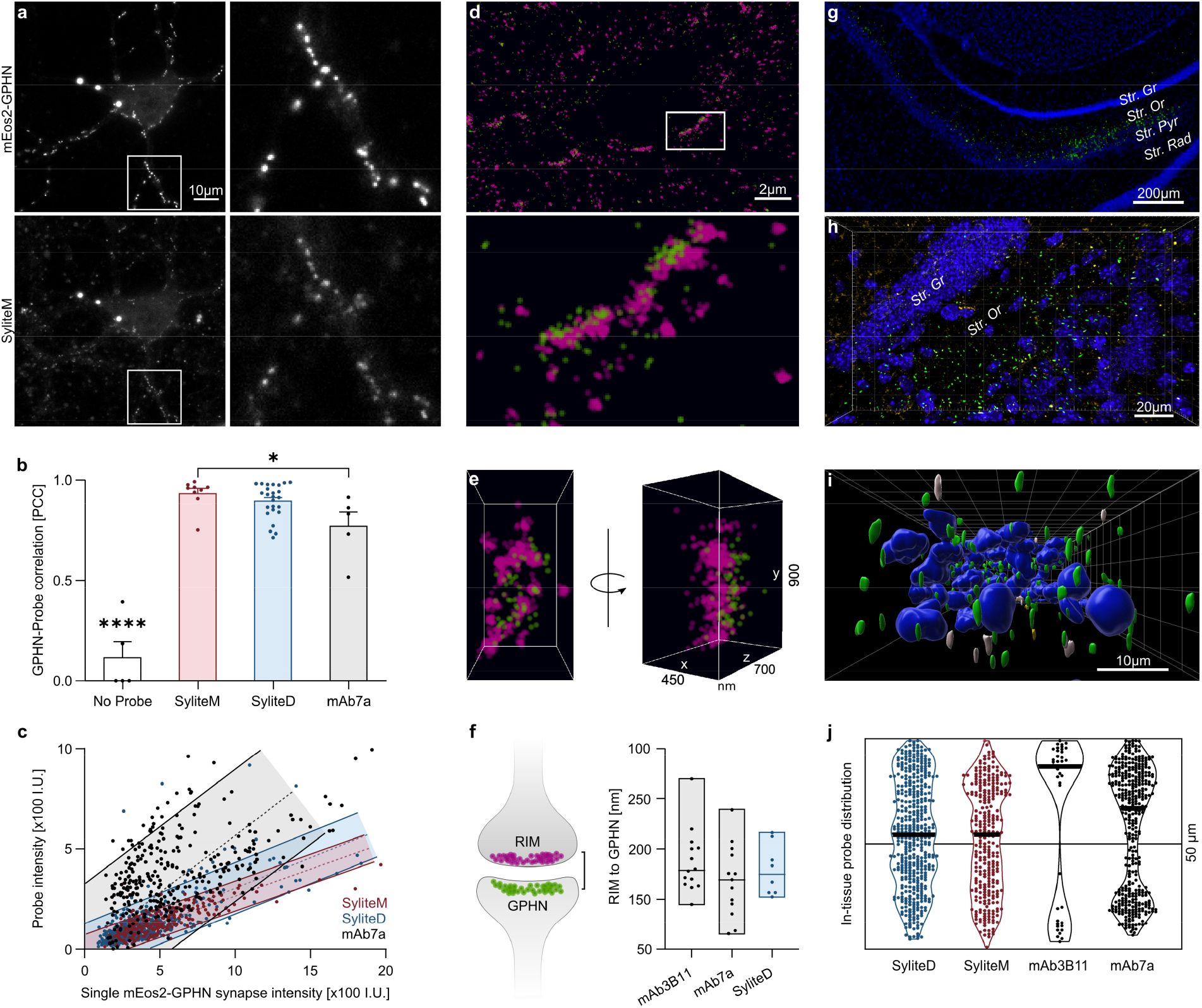
Sylites: versatile probes to visualize inhibitory synapses. **a-d. Sylites enable linear visualization of inhibitory synapses in neuronal cultures. a.** Top: Fixed cortical neurons expressing mEos2-gephyrin, bottom: SyliteM 50 nM staining of the fixed sample. Right: zoom in on the boxed region. SyliteM stains recombinant and endogenous gephyrin. **b.** Pearson’s correlation coefficients (PCC) of mEos2-gephyrin expressing neurons with the counterstain of SyliteM, SyliteD or mAb7a. All probes show high correlation to the recombinant neurons. Mean±SEM. Significance determined using one-way ANOVA with a follow up Tukey’s test for multiple comparisons. * P<0.05, **** P<0.0001. **c.** Intensity dependence of single mAb7a or Sylite labeled synapses to the reference mEos2-gephyrin synapse signal intensity. Much higher signal scattering is observed with mAb7a (grey), while both SyliteM (red) and SyliteD (blue) have a constant linear labeling behavior. Shaded regions indicate a 90% prediction interval. 10 pairs of images were used for each probe. **d-f. Super-resolution imaging and nanometric measurements with SyliteD. d.** Top: dSTORM of neuronal synapses with presynaptic RIM labeling with RIM1/2-CF680 antibody (magenta) and postsynaptic gephyrin labeling with SyliteD (green). Bottom: zoomed region. **e.** Side and *en face* view of a single synapse **f.** Nanometric distance measurement with SyliteD. RIM to gephyrin center of mass distance measurements conducted with RIM1/2-CF680 and either gephyrin antibodies or SyliteD. In all cases an average distance of ~130 nm was calculated. Bars indicate the full range of individual measurements, the in-bar line indicates the median. **g-j. Sylites reveal the distribution of inhibitory synapses in hippocampal sections. g.** Wide field 2D image of ventral hippocampus section stained with DAPI (blue) and SyliteD (green). Gephyrin staining is visible in the *stratum oriens* of the CA3 region of the ventral hippocampus. Str. Rad – Stratum Radiatum; Str. Pyr – Stratum Pyramidal; Str. Or – Stratum Oriens; Str. Gr – Stratum Granulosum. **h.** Confocal microscopy. 3D top view of SyliteD and mAb3B11 24-hour co-staining of ventral hippocampus section. Green – SyliteD, gold – mAb3B11, blue – DAPI nuclear staining. Numerous synapses are visible with SyliteD staining, mAb3B11 produces fewer detections. Synapses appear in the *stratum oriens.* **i.** 3D volumetric representation of nuclei and inhibitory synapses. Side view of a section co-labeled for gephyrin for 24 hours with SyliteD and mAb3B11. Green – SyliteD, gold – mAb3B11, blue – DAPI nuclear staining. In white SyliteD and mAb3B11 co-labeled synapses. **j.** Distribution of detected synapses in 50 μm-thick hippocampal sections after 24-hour staining. Violin plot represents the labeled synapse distribution. Thick black lines - median Z position of detected synapses. The hourglass shape of antibody labeling indicates skewed antibody distribution, towards the surfaces of sections.

Consequently, we conducted super-resolution nanoscopic distance measurements between the neuronal pre- and post-synapse using our probe and gephyrin antibodies. We carried out dSTORM experiments, focusing on the dimeric SyliteD due to its suitable blinking properties. Cortical neurons were labeled for gephyrin and RIM, a protein of the presynaptic active zone (AZ), using SyliteD together with primary anti-RIM1/2 and CF680-conjugated secondary antibodies (Fig.2d,e). Dual-color 3D-dSTORM images show that the SyliteD detections closely match the distribution of RIM in the AZ, confirming a recent finding of a strong association between RIM and gephyrin sub-synaptic domains^4^. The measured Euclidian distance between SyliteD and RIM1/2-CF680 was 129±24 nm (mean±SD), in agreement with the estimated molecular sizes separating the two labels^16^. The direct comparison with mAb7a and mAb3B11 labeling confirmed that SyliteD provides a precise read-out for the location of synaptic gephyrin and receptor binding sites at inhibitory synapses (Fig.2f).

Determining the organization, distribution and integrity of inhibitory synapses in brain tissue is of central importance for a wide range of neurobiological topics, from neural circuits to neuropathology. Until now, tissue staining of inhibitory synapses has been an elaborate and time-consuming procedure that was generally limited to relatively thin brain sections (≤ 16 μm) to obtain reliable labeling^7^. Here we demonstrate that Sylites, owing to their small size, effectively penetrate 50 μm-thick tissue sections, achieving high-contrast labeling within one hour, using a standard, immunohistochemistry protocol. We visualized inhibitory synapses and their distribution using epifluorescence microscopy with 20X magnification, giving us a macro-overview of the inhibitory synapse distribution in brain. (Fig.2g). Next, we incubated brain hippocampal sections for 1, 24 and 72 hours with Sylites and with mAb3B11 or mAb7a, then imaged the sections with a confocal microscope, deconvoluted the image stacks, and reconstructed 3D images. Sylite-visualized synapses were observed in the *stratum oriens* of the CA3 region of the ventral hippocampus, an area packed with inhibitory interneurons^17^ (Fig.2g,h, MovieS1). Sylites detected synaptic clusters throughout the entire section, demonstrating a complete penetration of the probe, already after 1 hour of incubation (MovieS2,3), and even after 72 hours of incubation with Sylites, we did not observe any significant background fluorescence (MovieS4,5). In contrast, after 24 hours, the antibody labeling was strongest near the surface of the section while the center remained largely unlabeled (Fig.2i-j, MovieS1,6). Antibody penetration improved after 72 hours; however, background staining was also higher (Fig.S7, MovieS4,5). This is seen by the drop in the overlap between antibody and Sylite labeling from ~0.4 for both 7a and 3B11 antibodies after 24h to ~0.1 after 72 hours (Fig.S7b). Lastly, 3D visualization of synapses produced by Sylites showed smooth, elongated and well-defined shapes of different sizes, in agreement with the known diversity of inhibitory synapses in the CNS^18^. After 24 hours, the antibodies produced both smooth and amorphous clusters, and after 72 hours, this pattern changed to primarily amorphous clusters and loss of any observable cluster directionality (MovieS7,8).

The past decades have seen a surge in technological advances in fluorescent microscopy and labeling methods, creating a demand for new probes, particularly for neurosciences, where micro- and nanoscale studies are required to decipher brain function^1^. However, only a handful of small affinity probes are available for fluorescence microscopy to date; the most prominent example is the widely used and easily applied DNA label DAPI. Sylites possess the same essential qualities as DAPI, namely high contrast visualization together with fast and reproducible staining protocols. Sylites bind neurotransmission-relevant gephyrin isoforms, acting as universal labels of inhibitory synapses, and their labeling linearity and defined stoichiometry enable deep quantification of the synapses. Our findings demonstrate that small affinity probes can be used adjuvant to antibodies or as their substitution, as they can be smartly designed to bind specific targets, and their small size effectively enables faster distribution, better penetration and staining in biological samples.

To summarize, our findings establish Sylites as powerful, versatile and reliable bioimaging tools for neuroscience. We anticipate that next-generation affinity probe development will continue to gather pace, as their synergy with cutting-edge microscopy is indisputable and will help to decipher brain cell organization and function.

## Acknowledgements

We thank Dr. Jens Vanselow, Stephanie Lamer and Alvaro Ciudad for supporting the mass spectrometric studies and analysis. We thank Prof. Eric Allemand and Dr. Fabrice Ango for providing gephyrin isoform constructs. And we thank Dr. Katharina Hemmen, Dr. Hanna Heil, Mike Friedrich and Jürgen Pinnecker for their guidance in 3D image analysis and light microscopy.

## Funding

H.M.M. acknowledges financial support from the Deutsche Forschungsgemeinschaft (DFG MA6957/1-1 and TRR 166 ReceptorLight, project B05).

## Author Contributions

Conceptualization, H.M.M, V.K., Methodology, H.M.M, C.G.S., V.K., C.S., S.L.R., O.A., C.V., K.H. A.S., O.S.F., P.T., Formal Analysis, H.M.M, C.G.S., V.K., C.S., S.L.R., O.A., A.S., O.S.F., P.T.; Investigation, H.M.M, C.G.S., V.K., C.S., S.L.R., O.S.F., R.W., N.F.N, S.K., C.V.; Writing – Original Draft Preparation, H.M.M. and V.K.; Writing – Review & Editing, H.M.M., V.K., C.G.S., C.S. and P.T. with help of co-authors; Visualization, H.M.M, C.G.S., V.K., C.S., S.L.R., O.A., A.S., P.T.; Supervision, H.M.M, C.G.S., C.V., K.H., A.S., O.S.F., P.T.; Project Administration, H.M.M.; Funding Acquisition, H.M.M.

## Competing interests

H. M.M. and V.K. filed a utility model concerning Sylites. C.Sc. is employed at Abbelight.

## Notes

### Summary of Updates

Supplementary Material added

## References

1. Choquet, D., Sainlos, M. & Sibarita, J. B. Advanced imaging and labelling methods to decipher brain cell organization and function. Nat. Rev. Neurosci. 22, 237–255 (2021).

2. Tyagarajan, S. K. & Fritschy, J.-M. Gephyrin: a master regulator of neuronal function? Nat. Rev. Neurosci. 15, 141–156 (2014).

3. Liu, Y. T. et al. Mesophasic organization of GABAA receptors in hippocampal inhibitory synapses. Nat. Neurosci. 23, 1589–1596 (2020).

4. Crosby, K. C. et al. Nanoscale Subsynaptic Domains Underlie the Organization of the Inhibitory Synapse. Cell Rep. 26, 3284–3297.e3 (2019).

5. Charrier, C. et al. A crosstalk between β1 and β3 integrins controls glycine receptor and gephyrin trafficking at synapses. Nat. Neurosci. 13, 1388–1395 (2010).

6. Gross, G. G. et al. An E3-ligase-based method for ablating inhibitory synapses. Nat. Methods 13, 673–678 (2016).

7. Schneider Gasser, E. M. et al. Immunofluorescence in brain sections: Simultaneous detection of presynaptic and postsynaptic proteins in identified neurons. Nat. Protoc. 1, 1887–1897 (2006).

8. Gross, G. G. et al. Recombinant Probes for Visualizing Endogenous Synaptic Proteins in Living Neurons. Neuron 78, 971–985 (2013).

9. Chamma, I. et al. Mapping the dynamics and nanoscale organization of synaptic adhesion proteins using monomeric streptavidin. Nat. Commun. 7, (2016).

10. Maric, H. M. et al. Gephyrin-binding peptides visualize postsynaptic sites and modulate neurotransmission. Nat. Chem. Biol. 13, 153–160 (2017).

11. Maric, H.-M., Mukherjee, J., Tretter, V., Moss, S. J. & Schindelin, H. Gephyrin-mediated γ-Aminobutyric Acid Type A and Glycine Receptor Clustering Relies on a Common Binding Site. J. Biol. Chem. 286, 42105–42114 (2011).

12. Maric, H. M. et al. Molecular basis of the alternative recruitment of GABAA versus glycine receptors through gephyrin. Nat. Commun. 5, 5767 (2014).

13. Fritschy, J. M., Harvey, R. J. & Schwarz, G. Gephyrin: where do we stand, where do we go? Trends Neurosci. 31, 257–264 (2008).

14. Kuhse, J. et al. Phosphorylation of gephyrin in hippocampal neurons by cyclin-dependent kinase CDK5 at Ser-270 is dependent on collybistin. J. Biol. Chem. 287, 30952–30966 (2012).

15. Specht, C. G. et al. Quantitative nanoscopy of inhibitory synapses: Counting gephyrin molecules and receptor binding sites. Neuron 79, 308–321 (2013).

16. Yang, X., Le Corronc, H., Triller, A. & Specht, C. G. Differential regulation of glycinergic and GABAergic nanocolumns at mixed inhibitory synapses. EMBO Rep. in press, (2021).

17. Cappaert, N. L. M., Van Strien, N. M. & Witter, M. P. The rat nervous system. (Academic Press, 2015). doi:10.1016/B978-0-12-374245-2.00020-6.

18. Santuy, A., Rodríguez, J. R., DeFelipe, J. & Merchán-Pérez, A. Study of the size and shape of synapses in the juvenile rat somatosensory cortex with 3D electron microscopy. eNeuro 5, (2018).

19. Prior, P. et al. Primary structure and alternative splice variants of gephyrin, a putative glycine receptor-tubulin linker protein. Neuron 8, 1161–1170 (1992).

