## Supplementary Material for "Sylites: Multipurpose markers for the visualization of inhibitory synapses"

**Outline**

### Methods

Unless otherwise noted, all resins and reagents were purchased from IRIS biotechnologies or Carl Roth and used without further purifications. All solvents used were HPLC grade. All water-sensitive reactions were performed in anhydrous solvents under positive pressure of argon. Statistical analysis was preformed using GraphPad Prism.

#### Peptide synthesis and fluorophore conjugation

Peptides were produced using standard solid phase peptide synthesis with Fmoc chemistry, shortly, 2-chlorotrityl resin (1.6 mmol/g) was swollen in dry Dichloromethane (DCM) for 30 min., then, the desired amino acid (AA) (1eq) and the Boc-Gly-OH (1eq) with 4 eq. of dry N,N-Diisopropylethylamine (DIEA) were added to the resin slurry. After overnight reaction at RT with agitation, the resin was capped with MeOH and washed with DCM and Dimethylformamide (DMF). Deprotection and conjugation cycles followed, where 20% piperidine solution in DMF was used to remove the Fmoc protecting group and after washes the peptide chain was elongated by adding AA (4 eq.) with Ethyl cyanohydroxyiminoacetate (Oxyma, 4 eq.) and N,N'-Diisopropylcarbodiimide (DIC, 4 eq.). Capping was done with DIEA (50 eq.) and acetic anhydride (50 eq.) in N-Methyl-2-pyrrolidone for 30 min. Coupling efficiency was monitored by measuring the absorption of the dibenzofulvene–piperidine adduct after deprotection. The peptides were cleaved from the resin using a cocktail of 90% TFA, 5% $H_2O$, 5% Triisopropylsilane, for 4 hours at RT. The peptides were precipitated in ice-cold ether and then purified with HPLC and analyzed by LC-MS.

The purified peptides or peptide dimers were conjugated with fluorophores either via $NH_2$-terminus using N-Hydroxysuccinimide (NHS) coupled dyes or via cysteine -SH side chain using maleimide coupled dyes. Shortly, for NHS coupling 1 eq. of peptide was dissolved in DMF with 3 eq. of DIEA and a fluorophore-NHS was added (1 eq for standard peptides, 2eq. for peptide dimers) and agitated overnight at 4°C. Maleimide conjugation was done with similar stoichiometry and conditions, but with 10 mM pH 7.4 PBS as a solvent and a minimal addition of DMSO to facilitate dissolution.

#### Peptide purification and characterization

The fluorescent probes were purified from the crude reaction mix by reverse phase HPLC using water acetonitrile gradient with 0.1% Formic Acid. LC-MS validation performed with similar gradient and LC-MS grade solvents. Semi-preparative HPLC was performed on Shimadzu Prominence equipped with a diode-array detector (DAD) system using a C18 reverse-phase column (Phenomenex Onyx Monolithic HD-C18 100 x 4.6 mm or Onyx Monolithic C18 100 x 10 mm). Purity and structural identity verified using a DAD equipped 1260 Infinity II HPLC with a C18 reverse-phase column (Onyx Monolithic C18 50 x 2 mm), coupled to a Mass Selective Detector single quadruple system (Agilent Technologies), ESI+ mode.

**Protein expression and purification**

Gephyrin P2 splice variant E domain (amino acids 318–736) was expressed in *E. Coli* and purified as described earlier[1]. Concisely, the protein was purified using via Intein-tag (Chitin beads, New England BioLabs), and after self-cleavage the protein was obtained by size-exclusion chromatography (SEC) column (HiLoad 16/600 Superdex 200pg, GE Healthcare) on an ÄKTA explorer system (GE Healthcare).

**Modeling of the SyliteD/gephyrin supracomplex**

Generation of the SyliteD bound to gephyrin E domain was carried out using the Rosetta FlexPepDock refinement protocol[2]. The crystal structure of gephyrin E domain bound to glycine receptor (GlyR) β subunit peptide (PDB ID: 4pd1) was used as a scaffold. Peptide residues where mutated using the Rosetta Fixed Backbone protocol to correspond to the binding sequence of SyliteD. Following this process, the linker and dye were added to demonstrate the feasibility of the dimer formation.

**Isothermal titration calorimetry (ITC)**

Measurements were performed using an ITC200 (MicroCal) at 25 °C and 1,000 revolutions per minute (rpm) stirring, PBS pH 7.4 was used as the standard solvent. Specifically, 40 µL of a solution 200 µM gephyrin E solution was titrated into the 200 µL sample cell containing 10 µM and 20 µM of SyliteD and SyliteM, respectively. In each experiment, a volume of 2.5 µL of ligand was added at a time resulting in 15 injections and a final molar ratio between 1:2 (SyliteM) and 1:4 (SyliteD). The dissociation constant ($K_D$) and stoichiometry (N) were obtained by data analysis using NITPIC, SEDPHAT and GUSSI[3]. Measurements were conducted three times for each probe and are given as mean values with the resulting standard deviations.

**Pulldowns with cellulose conjugated peptides**

Cellulose membrane bound peptides were produced using µSPOT solid phase peptide synthesis[1]. After completion of automated peptide synthesis, cellulose bound peptides side chains were deprotected with 90% TFA, 5% $H_2O$, 5% Triisopropylsilane for 3 hrs at RT, followed by washing with 5×2 mL $H_2O$. Afterwards, cellulose disks were left do dry overnight in a fume hood and stored at 4°C until use. For pulldowns, disks were first blocked in 2% (w/v) BSA in PBS for 1 hour at 25°C. Subsequently, one disc was incubated with 100 µL of mouse brain homogenate mixed with 100 µL of 10 mM TCEP in PBS for 45 min at 30 °C. After washing with 3×300 µL PBS, 50 µL loading buffer (NuPAGETM LDS-sample buffer, ThermoFisher Scientific) were added and incubation at 70°C for 2×5 min with a brief vortex in between followed. Samples were stored at -80°C until preparation for mass spectrometric proteomic analysis. The FSI**GV**SYPRRRRRRRRR, and (YSI**GV**SYPRpeg)$_2$KC non-binding analogues of SyliteM and SyliteD, respectively, were used for the assay. These sequences contain a binding-abolishing mutation, described in Maric et al. 2017[4].

**Mass spectrometric analysis of pulldowns**

Alkylation of the eluate was achieved by reduction with 50 mM dithiothreitol for 10 min at 70°C and 650 rpm in a thermoshaker followed by addition of 2-iodoacetamide to a final concentration of 120 mM and incubation in the dark for 20 min. Afterwards, cold acetone was added in a 4.5:1 ratio and overnight incubation at -20°C followed. Then, the samples were centrifuged at 12,000 g for 20 min at 4°C. Pellets were washed with 4×1 mL of cold acetone with 5 min centrifugations at 12,000 g in between. Next, pellets were left to dry under ventilation for 10 min. The protein pellet was resuspended in 50 µL of 8 M urea in 100 mM ammonium bicarbonate (ABC) using a bioruptor (diagenode) with 3 cycles for 30 sec. Afterwards, 50 µL of 100 mM ABC were added, followed by addition of 0.25 µg endoproteinase LysC. After incubation for 2 hrs in a thermoshaker at 30 °C and 900 rpm, 100 µL of 100 mM ABC and 0.25 µg trypsin were added. Following overnight incubation at 37 °C, the samples were acidified using 20 µL of 10% trifluoracetic acid (TFA). Stage tips were prepared by insertion of three C18 disks into a pipette tip. Each Stage tip was pre-washed with 50 µL MeOH, followed by 50 µL of 60% ACN with 0.3% (v/v) formic acid (FA), followed by equilibration with 2x50 µL of a 2% ACN solution with 0.3% (v/v) TFA. After sample loading, the tips were centrifuged for 10 min at 2,000 g and washed with 3×50 µL of 2% ACN with 0.3% (v/v) TFA. Elution was achieved using 2×50 µL of 60% ACN with 0.3% (v/v) FA, then the samples were lyophilized for storage until solubilization in 25 µL of 2% ACN with 0.1% (v/v) FA.

NanoLC-MS/MS analyses were performed on an Orbitrap Fusion (Thermo Scientific) equipped with a PicoView Ion Source (New Objective) and coupled to an EASY-nLC 1000 (Thermo Scientific). Peptides were loaded on capillary columns (PicoFrit, 30 cm x 150 µm ID, New Objective) self-packed with ReproSil-Pur 120 C18-AQ, 1.9 µm (Dr. Maisch) and separated with a 60-minute linear gradient from 3% to 30% acetonitrile and 0.1% formic acid and a flow rate of 500 nl/min.

Both MS and MS/MS scans were acquired in the Orbitrap analyzer with a resolution of 60,000 for MS scans and 7,500 for MS/MS scans. HCD fragmentation with 35 % normalized collision energy was applied. A Top Speed data-dependent MS/MS method with a fixed cycle time of 3 seconds was used. Dynamic exclusion was applied with a repeat count of 1 and an exclusion duration of 30 seconds; singly charged precursors were excluded from selection. Minimum signal threshold for precursor selection was set to 50,000. Predictive AGC was used with AGC a target value of 2E5 for MS scans and 5e4 for MS/MS scans. EASY-IC was used for internal calibration.

Raw MS data files were analyzed with MaxQuant version 1.6.2.2. Database search was performed with Andromeda, which is integrated in the utilized version of MaxQuant. The search was performed against the UniProt mus musculus reference proteome database (download date: 2020-08). Additionally, a database containing common contaminants was used. The search was performed with tryptic cleavage specificity with 3 allowed miscleavages. Protein identification was under control of the false-discovery rate (FDR; <1% FDR on protein and PSM level). In addition to MaxQuant default settings, the search was performed against following variable modifications: Protein N-terminal acetylation, Gln to pyro-Glu formation (N-term. Gln) and oxidation (Met). Carbamidomethyl (Cys) was set as fixed modification. Further data analysis was performed using R scripts developed in-house. LFQ intensities were used for protein quantitation. Proteins with less than two razor/unique peptides were removed. Missing LFQ intensities in the control samples were imputed with values close to the baseline. Data imputation was performed with values from a standard normal distribution with a mean of the 5% quantile of the combined log10-transformed LFQ intensities and a standard deviation of 0.1. For the identification of significantly enriched proteins, boxplot outliers were identified in intensity bins of at least 300 proteins. Log2 transformed protein ratios of sample versus control with values outside a 1.5x (significance 1) or 3x (significance 2) interquartile range (IQR), respectively, were considered as significantly enriched.

**HEK293 and COS7 cell culture**

HEK293 and COS7 cells were cultured in DMEM (GIBCO), supplemented with GlutaMax and pyruvate (GIBCO), 10% fetal bovine serum (GIBCO) and 1% Penicillin/Streptomycin (Sigma) at 37°C and with 5% $CO_2$. Stable HEK293 cells expressing eGFP-gephyrin were grown with 0.4 mg/mL of the selective antibiotic G418.

The cells were plated on 0.15 mm thick 18 mm glass coverslips (HEK293 on coverslips that were coated with 35 µg/ml Poly-D-Lysine) in a 12-well plate and were transfected with 1 µg plasmid DNA per coverslip using PEI (Polyethylenimine). The transfection was performed at 60-80% confluence. Shortly before transfection the medium was changed to fresh DMEM. The DNA was added to 100 µl DMEM without additives and mixed, 4 µl fresh PEI (1 mg/ml) was added, mixed immediately and incubated for 20 min at RT. The transfection mix was pipetted drop-wise on cells while swirling, and incubated overnight. The medium was changed to fresh DMEM with 2% FBS after 12-24 hours, and on the following day the cells were fixed and used for staining.

The following constructs were used for transient transfection of HEK293 and COS7 cells: eGFP-gephyrin P1[5] and eGFP - pEGFP-C2 were a gift from Prof. Matthias Kneussel (ZMNH, Germany); Venus-gephyrin[6] and pHluorin-tagged GlyR β-loop transmembrane protein[7] constructs (Specht lab); gephyrin isoform constructs (Supplementary Table 1 and Appendix 1) provided by Prof. Eric Allemand (INSERM, France) and Dr. Fabrice Ango (INSERM, France).

**Culture and infection of primary neurons**

Primary murine hippocampal neurons were prepared from wildtype CD-1 mice (Jackson Laboratory) at embryonic day 17 (E17). Experiments were approved by the local veterinary authority (Veterinäramt der Stadt Würzburg, Germany) and the Ethics Committee of Animal Experiments, Regierung von Unterfranken, Würzburg, Germany (FBVVL 568/200-324/13). Hippocampal neurons were grown in neurobasal medium (21103-049 Life Technologies, Massachusetts, USA) supplemented with 1% 200 mM L-Glutamine (25030-024 Life Technologies, Massachusetts, USA), 1% B27 (17504-044 Life Technologies, Massachusetts, USA). 50% of the medium was exchanged every 7 days in culture. 60,000 hippocampal neurons were seeded on 18 mm glass coverslips. Neurons were taken for experiments after three weeks in culture (day in vitro 21 = DIV21).

All procedures involving animals were in compliance with the regulations of the French Ministry of Agriculture and the Direction départementale des services vétérinaires de Paris (Ecole Normale Supérieure, animalerie des rongeurs, license B 75-05-20). Primary murine cortical neurons were dissociated from wildtype C57BL/6J mice (Janvier, France) at embryonic day 17 (E17) and cultured on 18 mm glass coverslips in neurobasal medium containing B27, glutamax and penicillin/streptomycin (all from Gibco). Where required, neurons were infected at day in vitro 1 to 5 (DIV1-5) with lentivirus driving the expression of full-length gephyrin tagged at its N-terminus with mEos2[8]. Neurons were used for experiments after two to three weeks in culture (DIV15-21).

**Cell fixation and immunocytochemistry**

Neurons, COS-7 and HEK293 cells were fixed in 0.1 M sodium phosphate buffer pH7.4 containing 4% paraformaldehyde (EM grade, Polysciences) and 1% sucrose for 10-20 min at 37°C. After three rinses in phosphate buffered saline (PBS), the cells were permeabilized in PBS containing 0.1% Triton X-100 for 10 min at room temperature, rinsed again and blocked for 1 h in PBS with 3% bovine serum albumin (BSA). Primary and secondary antibodies

were applied sequentially in blocking solution for 1 hour. The fluorescent probes were applied together with the primary antibody, unless otherwise mentioned.
Primary antibodies: mAb7a (147 011), mAb3B11 (147 111), Synaptic Systems.
Secondary antibodies were purchased from ThermoFisher: anti-mouse conjugated IgG with AlexaFluor (A) 647 (A-21235), A555 (A-21422), A488 (A-27023) or DyLight650 (84545).
Unless otherwise noted the Sylites were applied with 50 nM concentration, and both primary and secondary antibodies with 1:1000 concentration.

### Wide field fluorescence microscopy

Unless otherwise stated the coverslips with samples were inserted in an imaging chamber (Ludin Chamber Type 1, Life Imaging Services) and imaged in PBS. The measurements were taken from distinct samples with a sample size $\geq 2$, for each group. A series of images, used to generate the datapoints, were acquired from different regions of the sample, each region having a distinct group of cells.

### Probe profiling

SyliteD, SyliteM and TMR2i labeled COS-7 cells were imaged on an inverted Leica DMI6000B microscope with a 100x oil-immersion objective (NA 1.49) using a Leica DFC9000 GTC VSC-05760 sCMOS camera (16-bit, image pixel size: 130 nm). The following excitation and emission filters were chosen: excitation 470/40, emission 525/50 for gephyrin-eGFP and soluble eGFP; ex. 545/25, em. 605/70 for TMR2i (Tetramethylrhodamine); exc. 628/40, em. 692/40 for Sylites (Sulfo-Cyanine 5 – Cy5), 10 images were acquired at a frame rate (exposure time) of 100 ms and constant illumination intensity to ensure comparability. $n \geq 8$.

### Neuron imaging

Hippocampal neurons were imaged using the above-described setup. $n \geq 6$. Wide field imaging of cortical neurons was done on an inverted Nikon Eclipse Ti microscope with a 100x oil-immersion objective (NA 1.49) using an Andor iXon EMCCD camera (16-bit, image pixel size: 160 nm). The following excitation and emission filters were chosen: excitation 485/20, emission 525/30 for Alexa Fluor 488 and unconverted (green) mEos2; ex. 560/25, em. 607/36 for Cy3; exc. 650/13, em. 684/24 for Alexa Fluor 647 or Cy5 (SyliteD). Generally, 10 images were acquired at a frame rate (exposure time) of 100 ms and at variable illumination intensity using a mercury lamp (Intensilight, Nikon) and neutral density filters to maximize the signal while avoiding saturation. All images in one channel were taken with constant settings to ensure comparability. $n \geq 5$.

**2D image processing and analysis**

Image processing and analysis were carried out using Fiji[9] (**F**iji **I**s **J**ust **I**mageJ) with JACoP[10] (**J**ust **A**nother **Co**localization **P**lugin) plugin for colocalization analysis. Macros and scripts (Appendix 2) were written by V.K.

mEos2-gephyrin single synapse segmentation and intensity recording was done with Icy[11] 2.0.3.0 using "Wavelet Spot Detector" function in a custom protocol written by V.K. (Appendix 3). mEos2-gephyrin synaptic puncta were segmented, average intensity of individual punctum was determined and compared to the average intensity of the corresponding punctum in the far-red spectrum for either mAb7a with a secondary A647 antibody or the staining of Sylites.

**Dual-color dSTORM super-resolution imaging**

Neurons were fixed at DIV20 and immuno-labelled with primary rabbit anti-RIM1/2 antibody (Synaptic Systems, No. 140203, 1:250 dilution) and mouse anti-gephyrin (Synaptic Systems, mAb7a, No. 147011, 1:500) in blocking buffer for 2 hours. CF680-conjugated goat anti-rabbit secondary antibody (Biotium, No. 20818, one dye per IgG, 1:250) was co-applied with Alexa Fluor 647 (A647) – coupled donkey anti-mouse (1:500) or with the SyliteD probe at a final concentration of 500 nM for 2 h. Coverslips were mounted in dSTORM buffer (Abbelight SMART-kit) on cavity slides (Heinz Herenz, No 1042001), sealed with twinsil (Picodent) and imaged. The measurements were taken from distinct samples with a sample size ≥ 3, for each group.

All three fluorophores (Cy5, A647, CF680) photo-switch under reducing and oxygen-free buffer conditions, making them suitable for dSTORM single molecule imaging[12], which enables the localization of the emitters with sub-diffraction localization precision. Thanks to their close spectral proximity, Cy5 or A647 were excited and acquired simultaneously with CF680 in the same dSTORM buffer (Abbelight SMART-Kit) using a 640 nm laser (Oxxius), and their respective signals discriminated after single molecule localization using a spectral demixing strategy[13]. To implement spectral demixing dSTORM of SyliteD – (Cy5 or gephyrin-A647) and RIM1/2-CF680 we used a dual-view Abbelight SAFe360, equipped with two Hamamatsu Fusion sCMOS cameras and mounted on an Olympus Ix83 inverted microscope with a 100X 1.5NA TIRF objective. The SAFe360 uses astigmatic PSF engineering to extract the axial position and achieves quasi-isotropic 3D localization precision, and a long-pass dichroic mirror to split fluorescence from single emitters on the two cameras.

Single molecule localization, drift correction, spectral demixing, data visualization and cluster analysis[14] (DBSCAN) were performed with Abbelight NEO software, using a neighborhood radius eps = 150 nm and minPts = 50 minimum neighbors for the antibody labelling. To compensate for the lower number of detections generated by the SyliteD probe we adjusted the DBSCAN parameters to eps = 200 nm and minPts = 10. To measure the distance between presynaptic RIM and the postsynaptic gephyrin cluster, the centers of mass of the segmented clusters were determined in each fluorescence channels The Euclidean distance representing the average distance between the two-point clouds was then calculated for each cluster.

**Brain section preparation and staining**

Experiments were approved by the local veterinary authority (Veterinäramt der Stadt Würzburg, Germany) and the Ethics Committee of Animal Experiments, Regierung von Unterfranken, Würzburg, Germany (FBVVL 568/200-324/13). Wildtype C57BL/6J mice (Jackson Laboratory) were transcardially perfused via the left ventricle with ice-cold phosphate-buffer saline 1x (PBS1x) followed by ice-cold 4% paraformaldehyde (in PBS 1x). Brains were then removed, post-fixed in 4% PFA for 2 hours, cryoprotected in 30% sucrose/PBS for 48-72 hours and cut on a cryostat (Leica CM1950) in 50µm coronal slices. The immunohistochemistry was performed in free floating sections. Tissue sections were blocked with blocking solution (10% Donkey serum (Bio-rad) with 0.3% TritonX in PBS 1x) for 1 hour at RT, then fluorescent probes and primary antibodies were applied in blocking solution for 1 hour at RT, or 24/72 hours at 4°C. Then slices were washed 3 times with PBS and incubated with secondary antibody for 1 hour or 2 hours for the 24/72 hours staining protocol at RT. When no antibodies were applied the slices were incubated for 1h at RT with the probes. Labelled sections were then incubated with DAPI (1:5000) for 5 min at RT and washed again with PBS. Lastly, the sections were mounted onto a gelatin-coated slides using mowiol as the mounting medium. Following primary antibodies were used: gephyrin mouse mAb7a 1:1000 and mouse mAb3B11 1:1000. The fluorophore-tagged secondary antibody used was Alexa 555 donkey anti-mouse (1:1000).

**Wide field and confocal imaging of brain sections**

Wide-field 20x microscopy of brain sections was done with a Zeiss Axio Imager 2 equipped with a Plan-Apochromat 20x/0.8 M27 objective. Images were taken with an Axiocam 506 and pixel size of 0,454 x 0,454 µm. For excitation of DAPI a wavelength of 353 nm with LED-Module 385nm (power 6.08%) and for the probes a wavelength of 650nm with LED-Module 630nm (power 22.50%) was used. Emission wavelength for DAPI was 465nm and for the probes 673nm. Image acquisition was set using Zeiss Tiles module.

Labeled samples were imaged on a Leica SP8 (Leica) confocal microscope equipped with an HC PL APO CS2 63.0×/1.40-NA oil UV objective. Images were taken using a 200-Hz resonant scanner, 12-bits, a voxel size of 58 × 58 × 170 $nm^3$, a pinhole of 108.7 µm (1 AU). For excitation violet 405 nm LASOS diode laser (power 1-2%), yellow-green 561 nm DPSS laser (power 1-4%), red 633 nm HeNe laser (power 1-2%) were used. Emission light was registered with Leica PMT detectors set to the following spectral ranges: 415-465 nm, gain 750-850V (DAPI channel); 575-620 nm, gain 800-950V (Sulfo-Cyanine 3 channel); 645-700 nm, gain 750-850V (Cy5 channel). Image acquisition was performed in sequential frame scan mode, with concurrent 405 nm and 633 nm excitation and acquisition in corresponding ranges, followed by 561 nm excitation with an acquisition in Sulfo-Cyanine 3 channel. Bleaching was compensated with a linear gain increase of 30-40 V for an hour. The measurements were taken from distinct samples with a sample size of 4 for each group.

**3D image processing**

Confocal data was deconvoluted using a computed PSF (Huygens Professional package, Scientific Volume Imaging) with the following settings: Logarithmic vertical mapping function; 50-100 I.U. background estimation; max. 40 iterations, background to noise ratio of 5, 0.05 quality threshold, optimized iteration mode, auto brick layout. 3D and volumetric representation, segmentation and modeling of the deconvoluted images were done in Imaris (Oxford Instruments). The volumetric representation and segmentation were done with the following settings: Antibodies and Sylites (synapse segmentation): No smoothing, 10% intensity threshold, 800 Voxels volume threshold. DAPI (nuclei segmentation): 0.468 µm (8 pixel) smoothing, 5% intensity threshold, 10000 Voxels volume threshold.

The videos and section snapshots were created with Imaris (Oxford Instruments). Following settings for color intensity representation were used: DAPI 0-70% of the intensity dynamic range (0%-black, 70-100% - max intensity color); SyliteM/D 5-30% of the intensity dynamic range; mAb3B11 + secondary mouse A555 antibody 5-30% of the intensity dynamic range; mAb7a + secondary mouse A555 antibody 10-50% of the intensity dynamic range.

### Supplementary Discussion

#### Probe development

Earlier reported dimeric gephyrin E domain binders[15,16] displayed an exceptional affinity in low nanomolar range but required two purification and synthesis cycles. To improve yields and reduce costs we here use a double fmoc Lysin building block to dimerize directly on solid support. First, we designed and synthesized five tri-dioxaoctanoic acid dimerized peptides with different gephyrin binding sequence length and evaluated their binding to gephyrin E domain using isothermal titration calorimetry (Fig. 1S.a). The strongest binder had eight amino acid long binding sequence. Having determined the functionality of the Lysin-branched dimer we explored whether the linker type has an impact on the binding. To compare the different linker designs, we used the µSPOT[1] approach (a SPOT[17] and Celluspot[18] based peptide microarray synthesis method) for the comparison of eleven different dimeric linkers (Fig.S1b), all containing a core gephyrin binding sequence, and some containing previously described[4] affinity-enhancing mutations. Lastly, we tested how the length of the Gephyrin binding sequence influences the interaction, by truncating the eight amino acid sequence stepwise by one amino acid, to a minimum of three amino acids (Supplementary Table 2). The resulting 113 different dimeric binders and their monomeric counterparts were probed with gephyrin E domain in microarray format (Fig.S1b). We observed the most intense readouts for the binders having eight amino acids (Fig.S1c-d), in line with our ITC findings. Notably, binders with a modified core binding motif had higher intensity than the wildtype motif. Next, we analyzed what impact had the different linkers on the intensity readouts. For the wildtype sequence we saw an increase in intensity for linkers 01, 05, 06, 07, 08, 09, 10, 11 (Supplementary Table 4) while the monomeric binder and other dimeric binders had significantly lower intensity than the highest intensity binder FSIVGSLP10U (Fig.S1e). These differences could not be resolved with the mutated, higher-affinity binders, possibly due to on-array saturation of the binders with protein, leading to near-equal intensity readouts for these binders. Taking this into account we opted for dioxaoctanoic acid as a linker for the dimeric probe since it contributes several H-bridges, providing good water solubility and allowing effective and economic synthesis.

Using microarray-based assays we have recently defined the sequence requirements for the binding of native gephyrin, by probing gephyrin binding peptide microarrays with mouse brain homogenate[1] (Fig.S1f). Guided by our findings on optimal dimer architecture we here combined variants of this consensus binding motif and synthesized multiple different monovalent and dimeric peptides (Appendix 5) and conjugated them C- and N-terminally with sulfo-cyanine-5 (Cy5), Alexa Fluor 647 (A647) and rhodamine dyes (Supplementary Table 5). Cy5 and A647 are both suitable for dSTORM, with A647 being the brighter and more stable fluorophore[19], while silicon rhodamine (SiR) is STED compatible and was shown to work in live cell assays[20]. In addition, hydrophobicity and overall charges was adjusted via N-terminal elongation of the core binding motif or addition of Arginines (Fig 2s.a). Microscopy-based evaluation identified SyliteM and its dimeric counterpart SyliteD as fluorescent probes with best correlation, brightness and probe overlap (Fig 2S) as well as high target to off-target ratio (Fig 3S). Notably, compared to the previously reported gephyrin probe, TMR2i[4], Sylites show 10- and 150-fold improved contrast (Fig.1c, SyliteM and SyliteD, respectively) and are fully compatible with advanced super-resolution techniques like direct stochastic optical reconstruction (dSTORM, Fig.2.d-f).

### Supplementary Figures

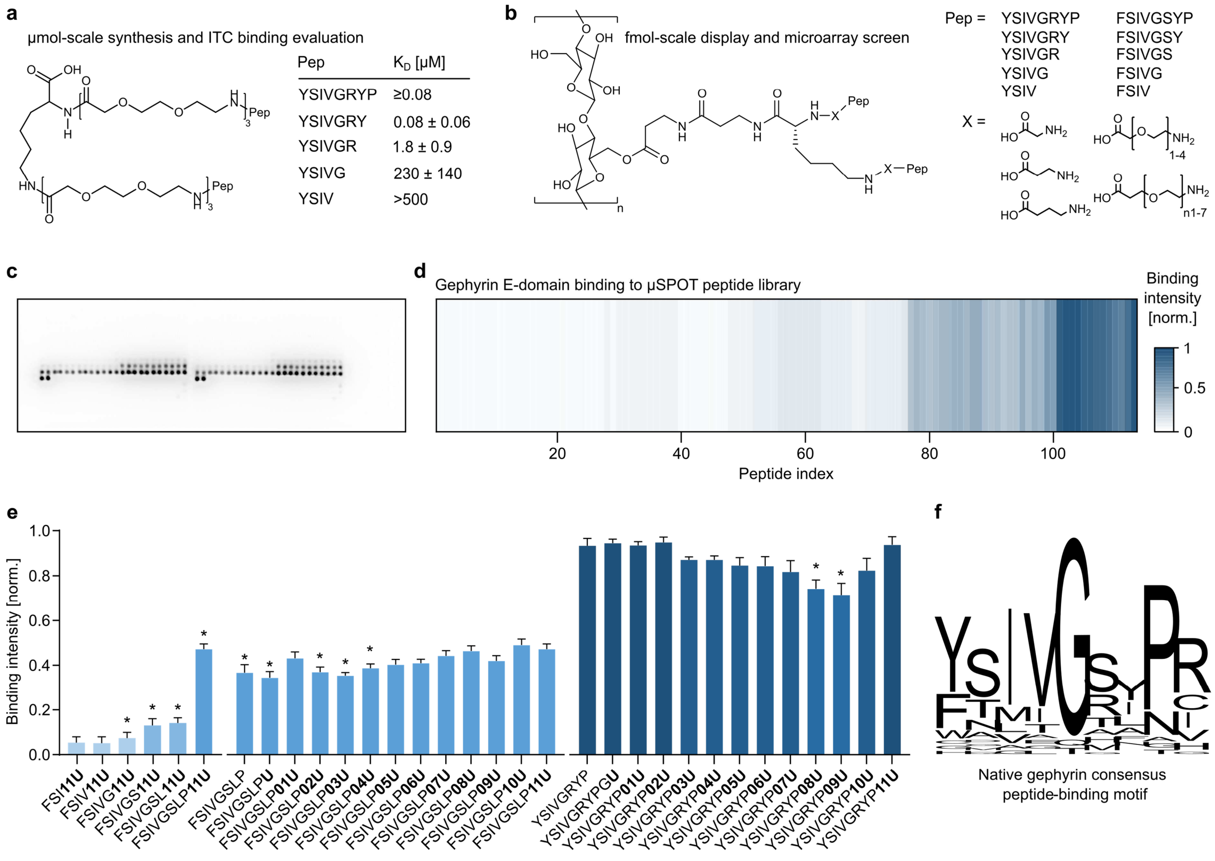

**Supplemental Figure 1. Optimization of probe sequence and multivalent architecture.** **a**. Validation of dimer design strategy via ITC. Gephyrin E affinity of dimers having different-length binding sequences was determined. Minimal binder sequence length is ≥5 amino acids. A ten-fold affinity increase is observed with the addition of, 7th and 8th amino acid to the binding sequence, the 8-mer binder having low-nanomolar affinity. **b.** µSPOT architecture of 113 monomeric and dimeric variants of gephyrin binding peptides. Peptides were based on ten binding sequences and eleven linker types (Supplementary Table 4). **c.** Representative example of the chemiluminescent gephyrin binding readout for the 113 monomeric and dimeric binders. Cellulose-conjugated peptides were printed on chips and incubated with 160pM of gephyrin E domain and subsequently detected with anti-gephyrin mAb3B11 and a secondary HRP conjugated antibody. Left and right are condition duplicates. **d**. Overview of the gephyrin binding intensities determined in microarray format. The intensities were internally normalized to the signal with highest intensity in the array and subsequently averaged and plotted for each peptide binder. **e.** The eight amino acid FSIVGSLP wild-type binding motif yields a higher luminescence signal than the shortened dimeric and monomeric analogues. The monomeric binder and **U**, **02U**, **03U**, **04U**, dimers have significantly lower intensity readout than the most intense wild-type binder FSIVGSLP**10U**. Monomer and dimers based on the enhanced binder sequence YSIVGRYP have the highest intensity readout in the microarray. **08U** and **09U** dimers have lower readouts than the most intense binder YSIVGRYP**11U**. One-way ANNOVA followed up by Tukey test for multiple comparisons was performed. Mean±SEM, $P<0.05$. n=6. **f.** Native gephyrin consensus peptide-binding motif determined by screening and alignment of fine mapped gephyrin-binding sites[1].

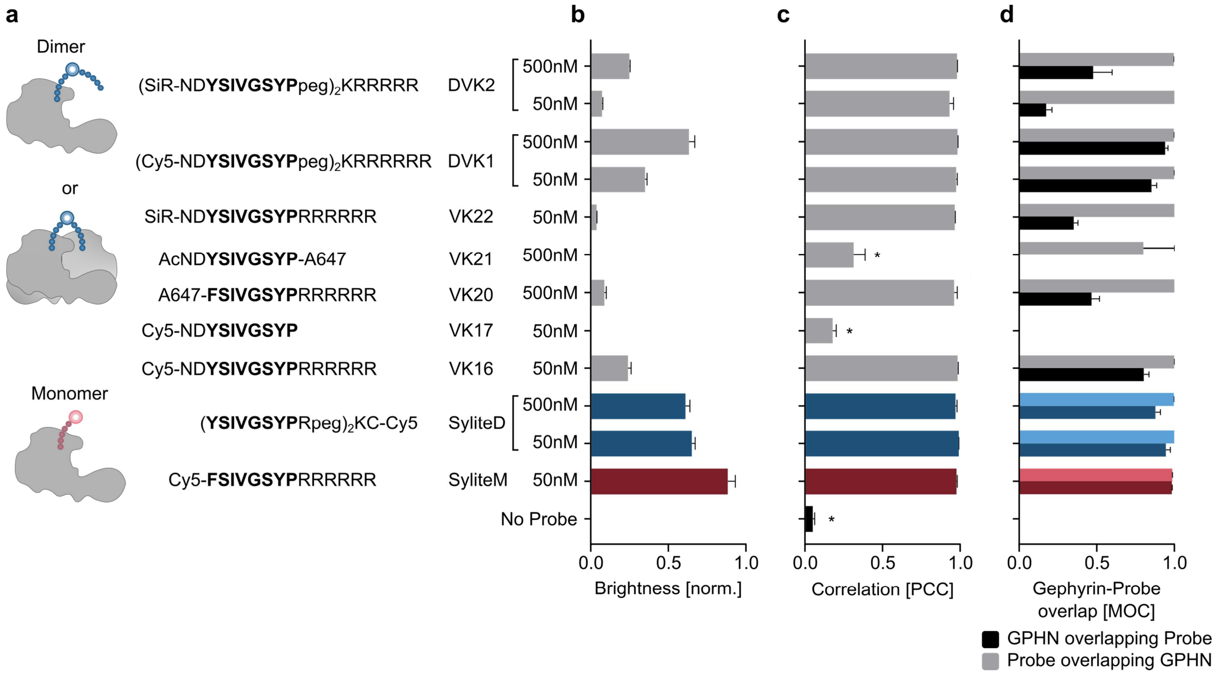

**Supplemental Figure 2. Microscopy-based probe evaluation.** Systematic comparison of the imaging properties of the synthesized fluorescent gephyrin probes in COS-7 cells expressing Venus-gephyrin. **a.** Binding modes of dimeric and monomeric probes together with sequences and fluorophores (Supplementary Table 5) of dimeric and monomeric probe variants (Appendix 5). The dimer can either bind one or two gephyrin molecules at once, while the monomer only one, always having stoichiometric 1:1 labeling. **b.** Comparison of the relative probe brightness. All probes are conjugated with far-red fluorophores, the average signal intensity coming from gephyrin clusters in far-red channel was divided by the corresponding average signal intensity of gephyrin clusters in green channel. Note that SyliteM is the brightest probe followed by SyliteD. **c.** Pearson's Correlation coefficients (PCC) of fluorescent probes and Venus-gephyrin. Next to the control only VK17 and VK21 show an incomplete correlation with gephyrin. **d.** Mander's overlap coefficients (MOC) - proportional coappearance of gephyrin and probe signals. When both values reach 1 it indicates an exclusive overlap of the two signals, meaning there is no over- or under-labeling of the target. Far-red signals coming from VK20, VK21, VK22 and DVK2 overlap almost completely with Venus-gephyrin, however Venus-gephyrin has only partial overlap with these probes, indicating under-labeling of the target. Significance was determined with ANOVA followed by Tukey's test for multiple comparisons. (*P<0.0001). Mean ± SEM.

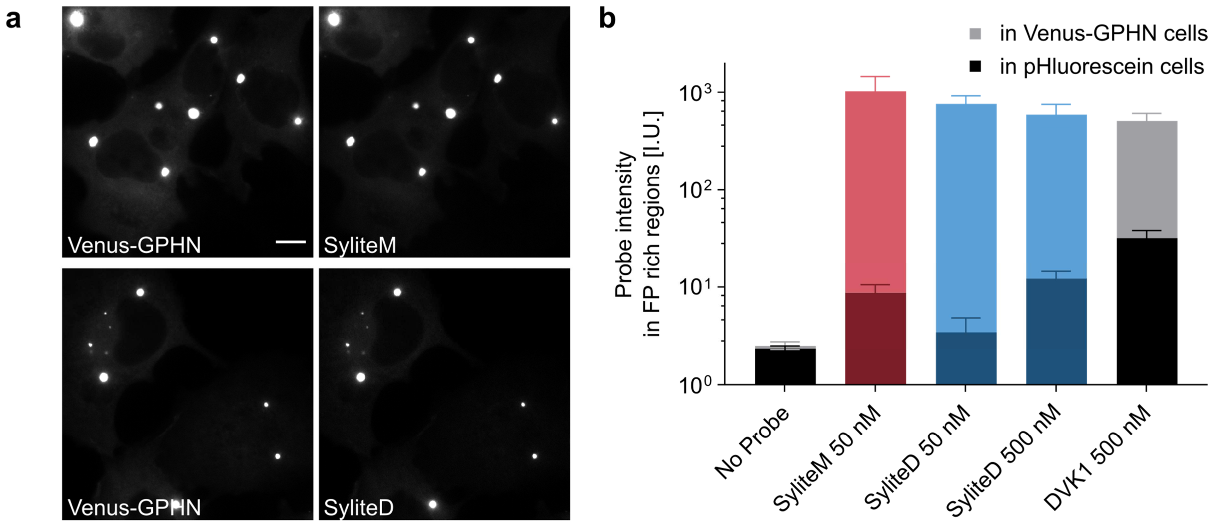

**Supplemental Figure 3. Sylites visualize gephyrin with high contrast. a**. Left: fixed COS7 cells expressing Venus-gephyrin. Right: 50 nM SyliteM and SyliteD staining of the fixed cells. Scale bar 10 µm. **b**. Target and off target labeling of the probes having the highest relative brightness, overlap and correlation to gephyrin. COS7 cells expressing pHluorescein-tagged GlyR β-loop transmembrane protein were used as control. Regions with fluorescent protein (FP) were segmented and their corresponding intensity in far-red (probe) channel was measured. Log scale of average on target and off target labeling intensities for the three probes. Target-to-off target staining ratios of ~120, ~220, ~50 and ~15 were calculated for SyliteM, SyliteD 50 and 500 nM, and DVK1 500 nM, respectively. Mean ± SD. n≥5.

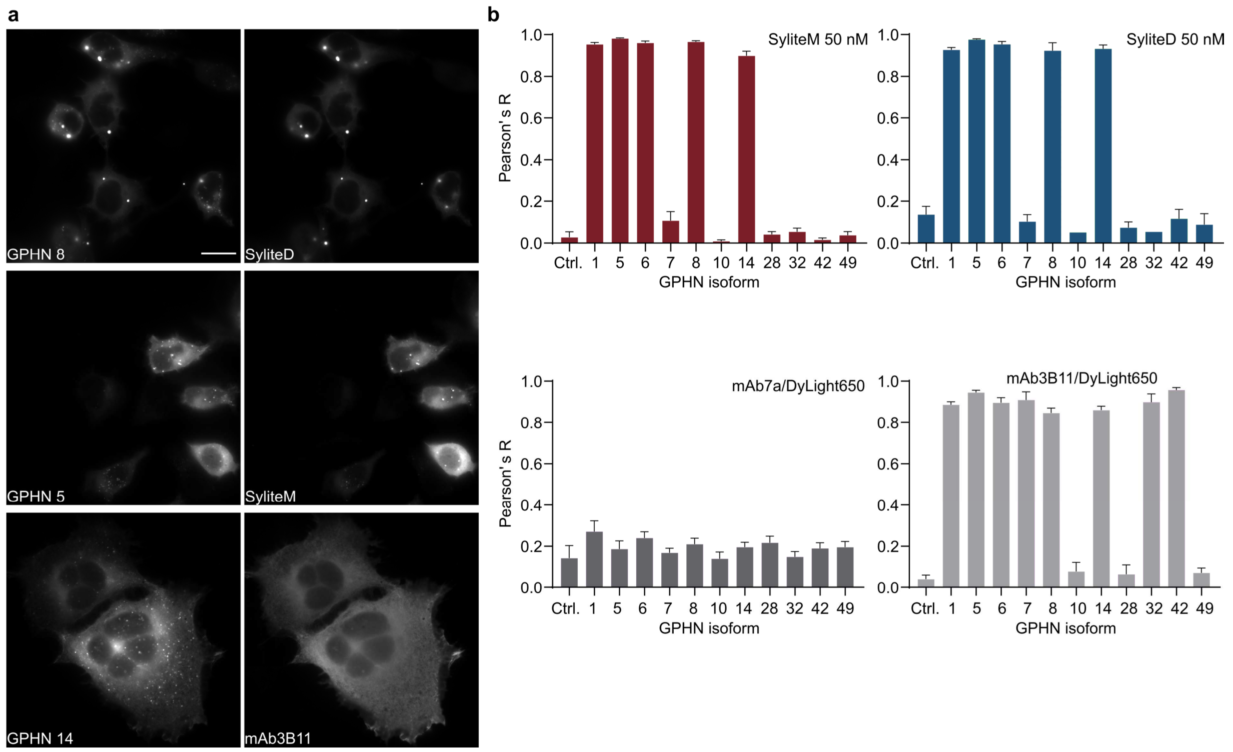

**Supplemental Figure 4. Isoform selectivity of Sylites and gephyrin antibodies. a.** Left HEK293 cells expressing mScarlet tagged gephyrin. Right: SyliteD, SyliteM (50 nM) and mAb3B11 with anti-mouse DyLight650 (1:1000) staining of the corresponding samples. Note that Sylites can label tightly packed gephyrin clusters, while mAb3B11 appears to label solely soluble gephyrin. **b.** Correlation between the fluorescent probes and gephyrin isoforms. The control group - HEK293 cells expressing mScarlet protein only. Average Pearson's R values (mean ± SEM). SyliteM and SyliteD significantly correlate with isoforms 1, 5, 6, 8, 14 (Supplementary Table 1). mAb3B11 correlates significantly with isoforms 1, 5, 6, 8, 14 and 7, 32, 42. Statistical significance determined with one-way ANOVA and a complimentary Dunnett's test for the comparison with a control group ($P<0.0001$). $n\geq5$.

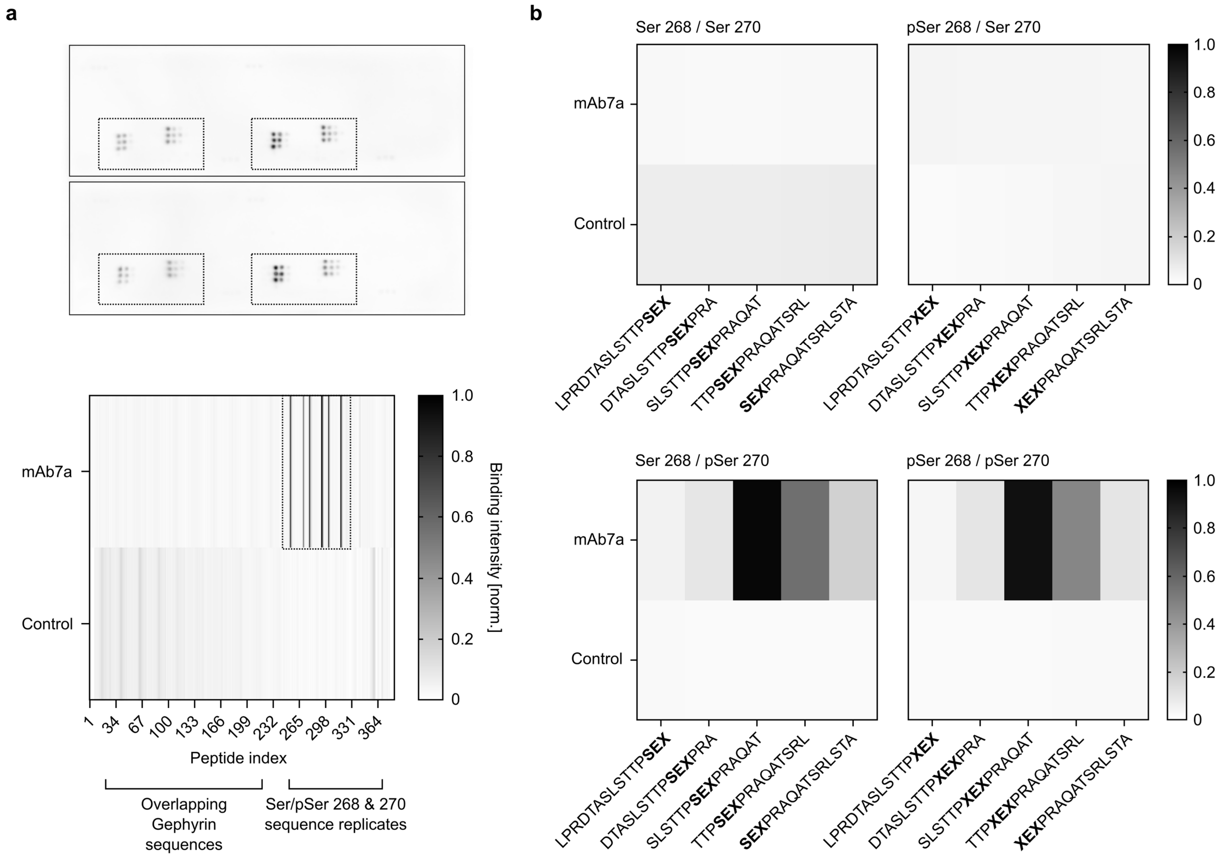

**Supplemental Figure 5. Gephyrin mAb7a explicitly binds a phosphorylated epitope. a.** Peptide microarray readout of overlapping gephyrin sequences. The gephyrin (GPHN-1 isoform)[21] sequence was displayed on the array with 15 amino-acid long peptides with 12 amino acid overlap between the peptides (Supplementary Table 3). mAb7a antibody was incubated on the microarray and detected with a secondary anti-mouse HRP conjugated antibody. Top panel: boxed - triplicates of phosphorylated peptide sequences. Bottom: a positional intensity readout, boxed is the region with phosphorylated sequence replicates. Intensities normalized to the highest intensity detected in the array. **b.** Averaged normalized intensity readout of the boxed region in a. X represents the phosphoserine. Chemiluminescent readout from SLSTTPSES$_{270}$PRAQAT is the most prominent, indicating mAb7a interaction with this linear epitope with phosphorylated Serin 270, phosphorylation of Ser 268 alone is not sufficient for antibody binding.

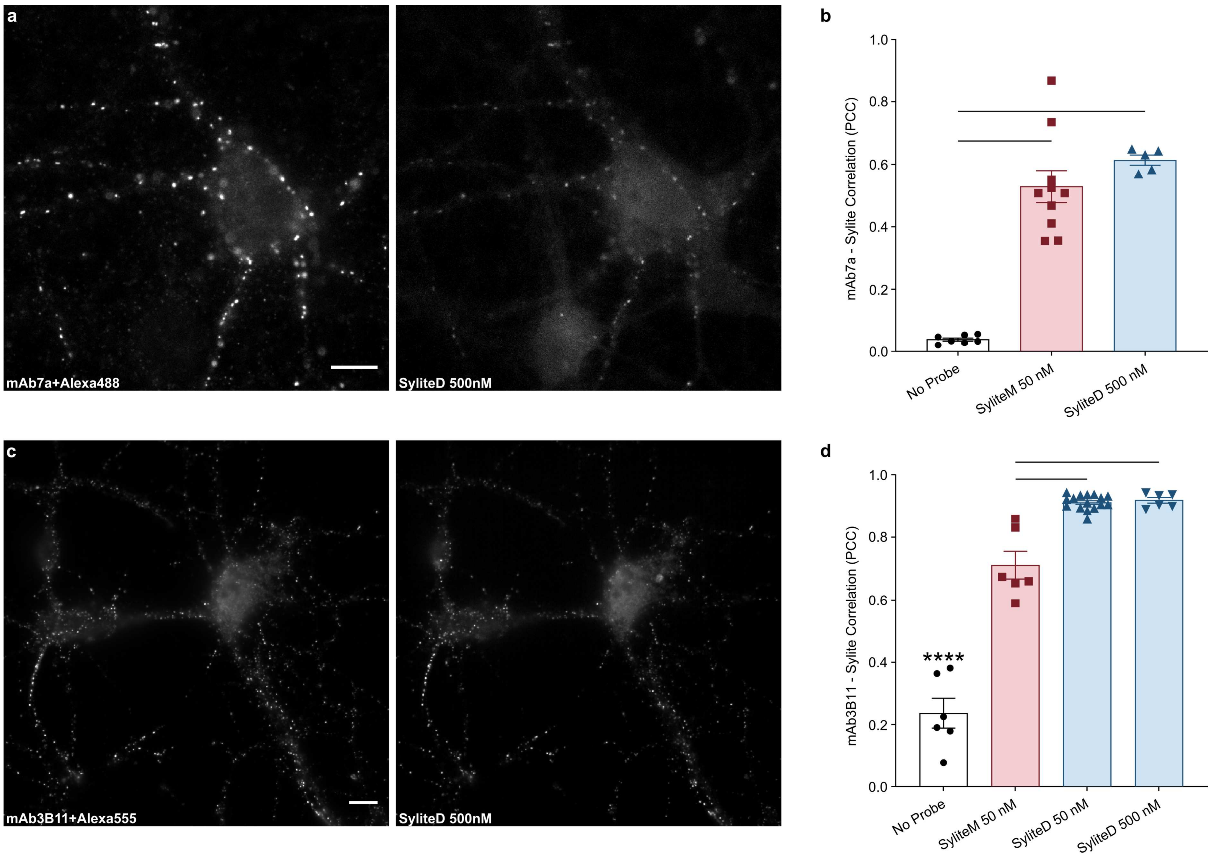

**Supplemental Figure 6. Sylites label inhibitory synapses in WT neurons. a.** DIV21 cortical neurons were fixed and co-stained with SyliteD and mAb7a. Similar pattern of antibody labeling and SyliteD is observable. Scale bar 10 µm. **b.** SyliteD and SyliteM staining correlates with the staining of mAb7a, but the degree of correlation is lower than with mAb3B11. These data are in-line with the narrow specificity of mAb7a, that labels only Ser270 phosphorylated neuronal gephyrin, therefore a lower correlation of mAb7a with Sylites is observed. On the “No Probe” group only a secondary mouse A488 antibody was applied. Mean ± SEM. Significance determined with one-way ANOVA combined with multiple comparison Tukey test. $P<0.0001$. **c.** DIV21 hippocampal neurons were fixed and co-stained with SyliteD and mAb3B11. Similar labeling pattern of mAb3B11 and SyliteD is seen. Scale bar 10 µm. **d.** SyliteD and mAb3b11 staining show significant and high correlation (PCC>0.91). Correlation of SyliteM is somewhat lower with PCC 0.71. On the “No Probe” group only secondary mouse A555 antibody was applied. Mean ± SEM. Significance determined with ANOVA combined with multiple comparison Tukey test. $P<0.0001$

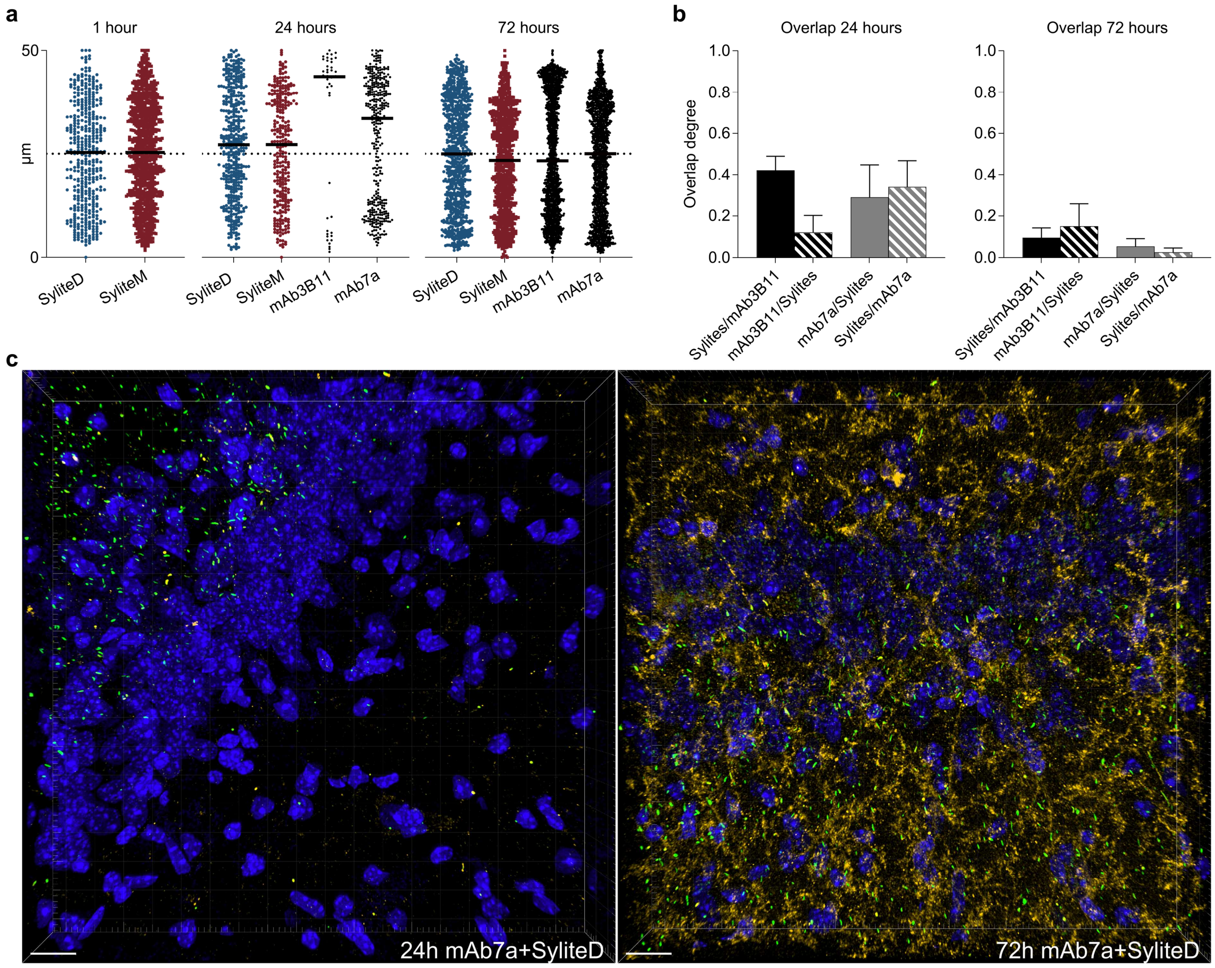

**Supplemental Figure 7. Sylites enable ultra-rapid synapse staining and visualization.** **a.** In-tissue distribution of gephyrin probes. A complete and homogenous penetration of Sylites is observed already after 1 hour of staining. Antibody penetration is inhomogeneous and incomplete after 24 hours, but better penetration is achieved after 72 hours. **b.** Sylite-antibody degree of overlap. 1 – full overlap, 0 – no overlap. After 24-hour staining of brain tissue about 40% of clusters detected by the antibodies are Sylite-positive as well. The number of clusters detected by Sylites is higher than the number of clusters detected with mAb3B11, hence only ~10% of Sylite-positive clusters are co-lableld with mAb3B11. After 72-hour staining the maximum degree of overlap drops to ~15%, in line with the increased unspecific staining of the antibodies. Mean ± SD. **c**. Sylites enable ultra-rapid synapse staining and visualization. Hippocampal section co-staining with SyliteD and mAb7a. Green – SyliteD, gold – mAb7a, blue – DAPI nuclear staining. Left – 24-hour staining. mAb7a and SyliteD clusters partially overlap. mAb7a appears to have some unspecific connective tissue staining, SyliteD synapse visualizations appear more consistent, a directional pattern of inhibitory synapse distribution can be observed. Right – 72-hour staining. Higher background staining with mAb7a is observed, the quality of SyliteD labeling does not change. Scale bar 15 µm.

### Supplementary Tables

**Supplementary Table 1. Gephyrin isoform constructs**

pLVX-hSyn-Flag-V5-mScarlett-IHRES-ZsGreen1 (version Eral)
pLVX-hSyn-Flag-V5-mScarlett-GPHN-1-IHRES-ZsGreen1 (version Eral)
pLVX-hSyn-Flag-V5-mScarlett-GPHN-5-IRES-ZsGreen1 (version Eral)
pLVX-hSyn-Flag-V5-mScarlett-GPHN-6-IRES-ZsGreen1 (version Eral)
pLVX-hSyn-Flag-V5-mScarlett-GPHN-7-IRES-ZsGreen1 (version Eral)
pLVX-hSyn-Flag-V5-mScarlett-GPHN-8-IRES-ZsGreen1 (version Eral)
pLVX-hSyn-Flag-V5-mScarlett-GPHN-10-IRES-ZsGreen1 (version Eral)
pLVX-hSyn-Flag-V5-mScarlett-GPHN-14-IRES-ZsGreen1 (version Eral)
pLVX-hSyn-Flag-V5-mScarlett-GPHN-28-IRES-ZsGreen1 (version Eral)
pLVX-hSyn-Flag-V5-mScarlett-GPHN-32-IRES-ZsGreen1 (version Eral)
pLVX-hSyn-Flag-V5-mScarlett-GPHN-42-IRES-ZsGreen1 (version Eral)
pLVX-hSyn-Flag-V5-mScarlett-GPHN-49-IRES-ZsGreen1 (version Eral)

**Supplementary Table 2. Peptide microarray 1 sequences**

| | | | |
|---|---|---|---|
| FSIG | FSIVG04K* | FSIVGS09K* | FSIVGSLP02G |
| FSIK* | FSIVG05K* | FSIVGS10K* | FSIVGSLP03G |
| FSI01K* | FSIVG06K* | FSIVGS11K* | FSIVGSLP04G |
| FSI02K* | FSIVG07K* | FSIVGSLG | FSIVGSLP05G |
| FSI03K* | FSIVG08K* | FSIVGSLK* | FSIVGSLP06G |
| FSI04K* | FSIVG09K* | FSIVGSL01K* | FSIVGSLP07G |
| FSI05K* | FSIVG10K* | FSIVGSL02K* | FSIVGSLP08G |
| FSI06K* | FSIVG11K* | FSIVGSL03K* | FSIVGSLP09G |
| FSI07K* | FSIVGG | FSIVGSL04K* | FSIVGSLP10G |
| FSI08K* | FSIVG01G | FSIVGSL05K* | FSIVGSLP11G |
| FSI09K* | FSIVG02G | FSIVGSL06K* | YSIVGRYPG |
| FSI10K* | FSIVG03G | FSIVGSL07K* | YSIVGRYPK* |
| FSI11K* | FSIVG04G | FSIVGSL08K* | YSIVGRYP01K* |
| FSIVG | FSIVG05G | FSIVGSL09K* | YSIVGRYP02K* |
| FSIVK* | FSIVG06G | FSIVGSL10K* | YSIVGRYP03K* |
| FSIV01K* | FSIVG07G | FSIVGSL11K* | YSIVGRYP04K* |
| FSIV02K* | FSIVG08G | FSIVGSLPK* | YSIVGRYP05K* |
| FSIV03K* | FSIVG09G | FSIVGSLP01K* | YSIVGRYP06K* |
| FSIV04K* | FSIVG10G | FSIVGSLP02K* | YSIVGRYP07K* |
| FSIV05K* | FSIVG11G | FSIVGSLP03K* | YSIVGRYP08K* |
| FSIV06K* | FSIVGSG | FSIVGSLP04K* | YSIVGRYP09K* |
| FSIV07K* | FSIVGSK* | FSIVGSLP05K* | YSIVGRYP10K* |
| FSIV08K* | FSIVGS01K* | FSIVGSLP06K* | YSIVGRYP11K* |
| FSIV09K* | FSIVGS02K* | FSIVGSLP07K* | |
| FSIV10K* | FSIVGS03K* | FSIVGSLP08K* | |
| FSIV11K* | FSIVGS04K* | FSIVGSLP09K* | |
| FSIVGK* | FSIVGS05K* | FSIVGSLP10K* | |
| FSIVG01K* | FSIVGS06K* | FSIVGSLP11K* | |
| FSIVG02K* | FSIVGS07K* | FSIVGSLPG | |
| FSIVG03K* | FSIVGS08K* | FSIVGSLP01G | |

### Supplementary Table 3. Peptide microarray 2 sequences

| | | | | |
|---|---|---|---|---|
| MATEGMILTNHDHQI | SKENILRASHSAVDI | IGHDIKRGECVLAKG | TTPXEXPRAQATSRL | TTPXEXPRAQATSRL |
| EGMILTNHDHQIRVG | NILRASHSAVDITKV | DIKRGECVLAKGTHM | XEXPRAQATSRLSTA | XEXPRAQATSRLSTA |
| ILTNHDHQIRVGVLT | RASHSAVDITKVARR | RGECVLAKGTHMGPS | - | - |
| NHDHQIRVGVLTVSD | HSAVDITKVARRHRM | CVLAKGTHMGPSEIG | - | - |
| HQIRVGVLTVSDSCF | VDITKVARRHRMSPF | AKGTHMGPSEIGLLA | - | - |
| RVGVLTVSDSCFRNL | TKVARRHRMSPFPLT | THMGPSEIGLLATVG | LPRDTASLSTTPSEX | LPRDTASLSTTPSEX |
| VLTVSDSCFRNLAED | ARRHRMSPFPLTSMD | GPSEIGLLATVGVTE | DTASLSTTPSEXPRA | DTASLSTTPSEXPRA |
| VSDSCFRNLAEDRSG | HRMSPFPLTSMDKAF | EIGLLATVGVTEVEV | SLSTTPSEXPRAQAT | SLSTTPSEXPRAQAT |
| SCFRNLAEDRSGINL | SPFPLTSMDKAFITV | LLATVGVTEVEVNKF | TTPSEXPRAQATSRL | TTPSEXPRAQATSRL |
| RNLAEDRSGINLKDL | PLTSMDKAFITVLEM | TVGVTEVEVNKFPVV | SEXPRAQATSRLSTA | SEXPRAQATSRLSTA |
| AEDRSGINLKDLVQD | SMDKAFITVLEMTPV | VTEVEVNKFPVVAVM | - | - |
| RSGINLKDLVQDPSL | KAFITVLEMTPVLGT | VEVNKFPVVAVMSTG | - | - |
| INLKDLVQDPSLLGG | ITVLEMTPVLGTEII | NKFPVVAVMSTGNEL | - | - |
| KDLVQDPSLLGGTIS | LEMTPVLGTEIINYR | PVVAVMSTGNELLNP | LPRDTASLSTTPXES | LPRDTASLSTTPXES |
| VQDPSLLGGTISAYK | TPVLGTEIINYRDGM | AVMSTGNELLNPEDD | DTASLSTTPXESPRA | DTASLSTTPXESPRA |
| PSLLGGTISAYKIVP | LGTEIINYRDGMGRV | STGNELLNPEDDLLP | SLSTTPXESPRAQAT | SLSTTPXESPRAQAT |
| LGGTISAYKIVPDEI | EIINYRDGMGRVLAQ | NELLNPEDDLLPGKI | TTPXESPRAQATSRL | TTPXESPRAQATSRL |
| TISAYKIVPDEIEEI | NYRDGMGRVLAQDVY | LNPEDDLLPGKIRDS | XESPRAQATSRLSTA | XESPRAQATSRLSTA |
| AYKIVPDEIEEIKET | DGMGRVLAQDVYAKD | EDDLLPGKIRDSNRS | - | - |
| IVPDEIEEIKETLID | GRVLAQDVYAKDNLP | LLPGKIRDSNRSTLL | - | - |
| DEIEEIKETLIDWCD | LAQDVYAKDNLPPFP | GKIRDSNRSTLLATI | - | - |
| EEIKETLIDWCDEKE | DVYAKDNLPPFPASV | RDSNRSTLLATIQEH | LPRDTASLSTTPXEX | LPRDTASLSTTPXEX |
| KETLIDWCDEKELNL | AKDNLPPFPASVKDG | NRSTLLATIQEHGYP | DTASLSTTPXEXPRA | DTASLSTTPXEXPRA |
| LIDWCDEKELNLILT | NLPPFPASVKDGYAV | TLLATIQEHGYPTIN | SLSTTPXEXPRAQAT | SLSTTPXEXPRAQAT |
| WCDEKELNLILTTGG | PFPASVKDGYAVRAA | ATIQEHGYPTINLGI | TTPXEXPRAQATSRL | TTPXEXPRAQATSRL |
| EKELNLILTTGGTGF | ASVKDGYAVRAADGP | QEHGYPTINLGIVGD | XEXPRAQATSRLSTA | XEXPRAQATSRLSTA |
| LNLILTTGGTGFAPR | KDGYAVRAADGPGDR | GYPTINLGIVGDNPD | - | - |
| ILTTGGTGFAPRDVT | YAVRAADGPGDRFII | TINLGIVGDNPDDLL | - | - |
| TGGTGFAPRDVTPEA | RAADGPGDRFIIGES | LGIVGDNPDDLLNAL | - | - |
| TGFAPRDVTPEATKE | DGPGDRFIIGESQAG | VGDNPDDLLNALNEG | LPRDTASLSTTPSEX | LPRDTASLSTTPSEX |
| APRDVTPEATKEVIE | GDRFIIGESQAGEQP | NPDDLLNALNEGISR | DTASLSTTPSEXPRA | DTASLSTTPSEXPRA |
| DVTPEATKEVIEREA | FIIGESQAGEQPTQT | DLLNALNEGISRANV | SLSTTPSEXPRAQAT | SLSTTPSEXPRAQAT |
| PEATKEVIEREAPGM | GESQAGEQPTQTVMP | NALNEGISRANVIIT | TTPSEXPRAQATSRL | TTPSEXPRAQATSRL |
| TKEVIEREAPGMALA | QAGEQPTQTVMPGQV | NEGISRANVIITSGG | SEXPRAQATSRLSTA | SEXPRAQATSRLSTA |
| VIEREAPGMALAMLM | EQPTQTVMPGQVMRV | ISRANVIITSGGVSM | - | - |
| REAPGMALAMLMGSL | TQTVMPGQVMRVTTG | ANVIITSGGVSMGEK | - | - |
| PGMALAMLMGSLNVT | VMPGQVMRVTTGAPI | IITSGGVSMGEKDYL | - | - |
| ALAMLMGSLNVTPLG | GQVMRVTTGAPIPCG | SGGVSMGEKDYLKQV | LPRDTASLSTTPXES | LPRDTASLSTTPXES |
| MLMGSLNVTPLGMLS | MRVTTGAPIPCGADA | VSMGEKDYLKQVLDI | DTASLSTTPXESPRA | DTASLSTTPXESPRA |
| GSLNVTPLGMLSRPV | TTGAPIPCGADAVVQ | GEKDYLKQVLDIDLH | SLSTTPXESPRAQAT | SLSTTPXESPRAQAT |
| NVTPLGMLSRPVCGI | APIPCGADAVVQVED | DYLKQVLDIDLHAQI | TTPXESPRAQATSRL | TTPXESPRAQATSRL |
| PLGMLSRPVCGIRGK | PCGADAVVQVEDTEL | KQVLDIDLHAQIHFG | XESPRAQATSRLSTA | XESPRAQATSRLSTA |
| MLSRPVCGIRGKTLI | ADAVVQVEDTELIRE | LDIDLHAQIHFGRVF | - | - |
| RPVCGIRGKTLIINL | VVQVEDTELIRESDD | DLHAQIHFGRVFMKP | - | - |
| CGIRGKTLIINLPGS | VEDTELIRESDDGTE | AQIHFGRVFMKPGLP | - | - |
| RGKTLIINLPGSKKG | TELIRESDDGTEELE | HFGRVFMKPGLPTTF | LPRDTASLSTTPXEX | LPRDTASLSTTPXEX |
| TLIINLPGSKKGSQE | IRESDDGTEELEVRI | RVFMKPGLPTTFATL | DTASLSTTPXEXPRA | DTASLSTTPXEXPRA |
| INLPGSKKGSQECFQ | SDDGTEELEVRILVQ | MKPGLPTTFATLDID | SLSTTPXEXPRAQAT | SLSTTPXEXPRAQAT |
| PGSKKGSQECFQFIL | GTEELEVRILVQARP | GLPTTFATLDIDGVR | TTPXEXPRAQATSRL | TTPXEXPRAQATSRL |
| KKGSQECFQFILPAL | ELEVRILVQARPGQD | TTFATLDIDGVRKII | XEXPRAQATSRLSTA | XEXPRAQATSRLSTA |
| SQECFQFILPALPHA | VRILVQARPGQDIRP | ATLDIDGVRKIIFAL | | |
| CFQFILPALPHAIDL | LVQARPGQDIRPIGH | DIDGVRKIIFALPGN | | |
| FILPALPHAIDLLRD | ARPGQDIRPIGHDIK | GVRKIIFALPGNPVS | | |
| PALPHAIDLLRDAIV | GQDIRPIGHDIKRGE | KIIFALPGNPVSAVV | | |
| PHAIDLLRDAIVKVK | IRPIGHDIKRGECVL | FALPGNPVSAVVTCN | | |
| IDLLRDAIVKVKEVH | IGHDIKRGECVLAKG | PGNPVSAVVTCNLFV | | |
| LRDAIVKVKEVHDEL | DIKRGECVLAKGTHM | PVSAVVTCNLFVVPA | | |
| AIVKVKEVHDELEDL | RGECVLAKGTHMGPS | AVVTCNLFVVPALRK | | |
| KVKEVHDELEDLPSP | CVLAKGTHMGPSEIG | TCNLFVVPALRKMQG | | |
| EVHDELEDLPSPPPP | AKGTHMGPSEIGLLA | LFVVPALRKMQGILD | | |
| DELEDLPSPPPPLSP | THMGPSEIGLLATVG | VPALRKMQGILDPRP | | |
| EDLPSPPPPLSPPPT | GPSEIGLLATVGVTE | LRKMQGILDPRPTII | | |
| PSPPPPLSPPPTTSP | EIGLLATVGVTEVEV | MQGILDPRPTIIKAR | | |
| PPPLSPPPTTSPHKQ | LLATVGVTEVEVNKF | ILDPRPTIIKARLSC | | |
| LSPPPTTSPHKQTED | TVGVTEVEVNKFPVV | PRPTIIKARLSCDVK | | |
| PPTTSPHKQTEDKGV | VTEVEVNKFPVVAVM | TIIKARLSCDVKLDP | | |
| TSPHKQTEDKGVQCE | VEVNKFPVVAVMSTG | KARLSCDVKLDPRPE | | |
| HKQTEDKGVQCEEEE | NKFPVVAVMSTGNEL | LSCDVKLDPRPEYHR | | |
| TEDKGVQCEEEEEEK | PVVAVMSTGNELLNP | DVKLDPRPEYHRCIL | | |
| KGVQCEEEEEEKKDS | AVMSTGNELLNPEDD | LDPRPEYHRCILTWH | | |
| QCEEEEEEKKDSGVA | STGNELLNPEDDLLP | RPEYHRCILTWHHQE | | |
| EEEEEKKDSGVASTE | NELLNPEDDLLPGKI | YHRCILTWHHQEPLP | | |
| EEKKDSGVASTEDSS | LNPEDDLLPGKIRDS | CILTWHHQEPLPWAQ | | |
| KDSGVASTEDSSSSH | EDDLLPGKIRDSNRS | TWHHQEPLPWAQSTG | | |

| | | |
|---|---|---|
| GVASTEDSSSSHITA | LLPGKIRDSNRSTLL | HQEPLPWAQSTGNQM |
| STEDSSSSHITAAAL | GKIRDSNRSTLLATI | PLPWAQSTGNQMSSR |
| DSSSSHITAAALAAK | RDSNRSTLLATIQEH | WAQSTGNQMSSRLMS |
| SSHITAAALAAKIPD | NRSTLLATIQEHGYP | STGNQMSSRLMSMRS |
| ITAAALAAKIPDSII | TLLATIQEHGYPTIN | NQMSSRLMSMRSANG |
| AALAAKIPDSIISRG | ATIQEHGYPTINLGI | SSRLMSMRSANGLLM |
| AAKIPDSIISRGVQV | QEHGYPTINLGIVGD | LMSMRSANGLLMLPP |
| IPDSIISRGVQVLPR | GYPTINLGIVGDNPD | MRSANGLLMLPPKTE |
| SIISRGVQVLPRDTA | TINLGIVGDNPDDLL | ANGLLMLPPKTEQYV |
| SRGVQVLPRDTASLS | LGIVGDNPDDLLNAL | LLMLPPKTEQYVELH |
| VQVLPRDTASLSTTP | VGDNPDDLLNALNEG | LPPKTEQYVELHKGE |
| LPRDTASLSTTPSES | NPDDLLNALNEGISR | KTEQYVELHKGEVVD |
| DTASLSTTPSESPRA | DLLNALNEGISRANV | QYVELHKGEVVDVMV |
| SLSTTPSESPRAQAT | NALNEGISRANVIIT | ELHKGEVVDVMVIGRL |
| TTPSESPRAQATSRL | NEGISRANVIITSGG | - |
| SESPRAQATSRLSTA | ISRANVIITSGGVSM | - |
| PRAQATSRLSTASCP | ANVIITSGGVSMGEK | - |
| QATSRLSTASCPTPK | IITSGGVSMGEKDYL | LPRDTASLSTTPSEX |
| SRLSTASCPTPKVQS | SGGVSMGEKDYLKQV | DTASLSTTPSEXPRA |
| STASCPTPKVQSRCS | VSMGEKDYLKQVLDI | SLSTTPSEXPRAQAT |
| SCPTPKVQSRCSSKE | GEKDYLKQVLDIDLH | TTPSEXPRAQATSRL |
| TPKVQSRCSSKENIL | DYLKQVLDIDLHAQI | SEXPRAQATSRLSTA |
| VQSRCSSKENILRAS | KQVLDIDLHAQIHFG | - |
| RCSSKENILRASHSA | LDIDLHAQIHFGRVF | - |
| | DLHAQIHFGRVFMKP | - |
| | AQIHFGRVFMKPGLP | LPRDTASLSTTPXES |
| | HFGRVFMKPGLPTTF | DTASLSTTPXESPRA |
| | RVFMKPGLPTTFATL | SLSTTPXESPRAQAT |
| | MKPGLPTTFATLDID | TTPXESPRAQATSRL |
| | GLPTTFATLDIDGVR | XESPRAQATSRLSTA |
| | TTFATLDIDGVRKII | - |
| | ATLDIDGVRKIIFAL | - |
| | DIDGVRKIIFALPGN | - |
| | GVRKIIFALPGNPVS | LPRDTASLSTTPXEX |
| | KIIFALPGNPVSAVV | DTASLSTTPXEXPRA |
| | FALPGNPVSAVVTCN | SLSTTPXEXPRAQAT |

520

521

**Supplementary Table 4. Linker building blocks**

| Abbreviation | Name |
|---|---|
| 01 | Glycine (G) |
| 02 | β-Alanine (A*) |
| 03 | GABA (GABA) |
| 04 | Oxapentanoic acid (O1P) |
| 05 | Amino-PEG1 acid (Peg1) |
| 06 | Dioxaoctanoic acid (O2Oc) |
| 07 | Amino-PEG2-acid (Peg2) |
| 08 | Amino-PEG3-Formicacid (Peg3) |
| 09 | Amino-PEG4-Formicacid (Peg4) |
| 10 | Amino-PEG5-acid (Peg5) |
| 11 | Amino-PEG7-acid (Peg7) |

**Supplementary Table 5. Fluorescent dyes**

| Abbreviation | Name | Structure |
| --- | --- | --- |
| Cy5 | Sulfo-Cyanine 5 | 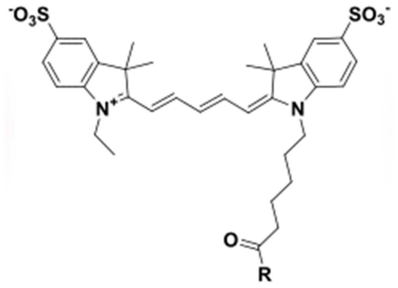 |
| A647 | Alexa647 | 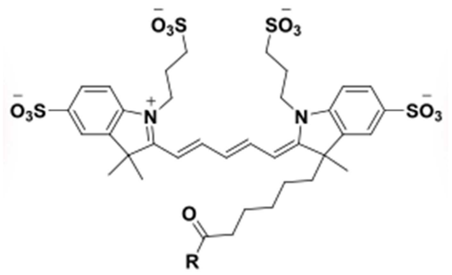 |
| SiR | Silicone Rhodamine | 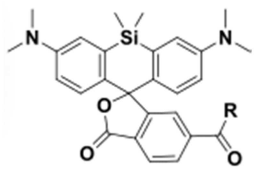 |
| TMR | TAMRA (Carboxytetramethylrhodamine) | 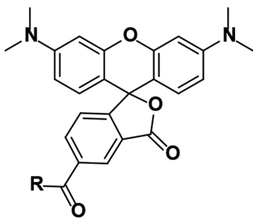 |

### Supplementary References

1. Schulte, C. *et al.* High-throughput Determination of Protein Affinities using Unmodified Peptide Libraries in Nanomolar Scale. *iScience* 101898 (2020).
2. Raveh, B., London, N. & Schueler-Furman, O. Sub-angstrom modeling of complexes between flexible peptides and globular proteins. *Proteins Struct. Funct. Bioinforma.* **78**, 2029–2040 (2010).
3. Brautigam, C. A., Zhao, H., Vargas, C., Keller, S. & Schuck, P. Integration and global analysis of isothermal titration calorimetry data for studying macromolecular interactions. *Nat. Protoc.* **11**, 882–894 (2016).
4. Maric, H. M. *et al.* Gephyrin-binding peptides visualize postsynaptic sites and modulate neurotransmission. *Nat. Chem. Biol.* **13**, 153–160 (2017).
5. Tyagarajan, S. K. *et al.* Extracellular signal-regulated kinase and glycogen synthase kinase 3β regulate gephyrin postsynaptic aggregation and GABAergic synaptic function in a calpain-dependent mechanism. *J. Biol. Chem.* **288**, 9634–9647 (2013).
6. Hanus, C., Ehrensperger, M. V. & Triller, A. Activity-dependent movements of postsynaptic scaffolds at inhibitory synapses. *J. Neurosci.* **26**, 4586–4595 (2006).
7. Specht, C. G. *et al.* Regulation of glycine receptor diffusion properties and gephyrin interactions by protein kinase C. *EMBO J.* **30**, 3842–3853 (2011).
8. Patrizio, A., Renner, M., Pizzarelli, R., Triller, A. & Specht, C. G. Alpha subunit-dependent glycine receptor clustering and regulation of synaptic receptor numbers. *Sci. Rep.* **7**, 1–11 (2017).
9. Schindelin, J. *et al.* Fiji: an open-source platform for biological-image analysis. *Nat. Methods* **9**, 676–682 (2012).
10. Bolte, S. & Cordelières, F. P. A guided tour into subcellular colocalization analysis in light microscopy. *J. Microsc.* **224**, 213–232 (2006).
11. De Chaumont, F. *et al.* Icy: An open bioimage informatics platform for extended reproducible research. *Nat. Methods* **9**, 690–696 (2012).
12. Heilemann, M. *et al.* Subdiffraction-resolution fluorescence imaging with conventional fluorescent probes. *Angew. Chemie - Int. Ed.* **47**, 6172–6176 (2008).
13. Lampe, A., Haucke, V., Sigrist, S. J., Heilemann, M. & Schmoranzer, J. Multi-colour direct STORM with red emitting carbocyanines. *Biol. Cell* **104**, 229–237 (2012).
14. Ester, M., Kriegel, H.-P., Sander, J., Xu, X. & others. A density-based algorithm for discovering clusters in large spatial databases with noise. in *Kdd* vol. 96 226–231 (1996).
15. Maric, H. M., Kasaragod, V. B. & Schindelin, H. Modulation of gephyrin-glycine receptor affinity by multivalency. *ACS Chem. Biol.* **9**, 2554–2562 (2014).
16. Maric, H. M. *et al.* Design and Synthesis of High-Affinity Dimeric Inhibitors Targeting the Interactions between Gephyrin and Inhibitory Neurotransmitter Receptors. *Angew. Chemie Int. Ed.* **54**, (2014).
17. Frank, R. Spot-synthesis: an easy technique for the positionally addressable, parallel chemical synthesis on a membrane support. *Tetrahedron* **48**, 9217–9232 (1992).
18. Dikmans, A., Beutling, U., Schmeisser, E., Thiele, S. & Frank, R. SC2: A novel process for manufacturing multipurpose high-density chemical microarrays. *QSAR Comb. Sci.* **25**, 1069–1080 (2006).
19. Helmerich, D. A., Beliu, G., Matikonda, S. S., Schnermann, M. J. & Sauer, M. Photoblueing of organic dyes can cause artifacts in super-resolution microscopy. *Nat. Methods* **18**, 253–257 (2021).
20. Lukinavičius, G. *et al.* Fluorogenic probes for live-cell imaging of the cytoskeleton. *Nat. Methods* **11**, 731–733 (2014).
21. Prior, P. *et al.* Primary structure and alternative splice variants of gephyrin, a putative glycine receptor-tubulin linker protein. *Neuron* **8**, 1161–1170 (1992).

### Supplementary Notes

#### Supplemetary Method 1. Imaging-based screening

Wide field imaging of labelled cells was done on an inverted Nikon Eclipse Ti microscope with a 100x oil-immersion objective (NA 1.49) using an Andor iXon EMCCD camera (16-bit, image pixel size: 160 nm). The following excitation and emission filters were chosen: excitation 485/20, emission 525/30 for Alexa Fluor 488 and unconverted (green) mEos2; ex. 560/25, em. 607/36 for Cy3; exc. 650/13, em. 684/24 for Alexa Fluor 647 or Cy5 (SyliteD). Generally, 10 images were acquired at a frame rate (exposure time) of 100 ms and at variable illumination intensity using a mercury lamp (Intensilight, Nikon) and neutral density filters to maximize the signal while avoiding saturation. All images in one channel were taken with constant settings to ensure comparability.

#### Supplemetary Method 2. Peptide microarray synthesis

μSPOT [1] peptide microarrays were synthesized using a Celluspot-based approach[18] using a MultiPep RSi robot (CEM GmbH) on in-house produced, acid labile, amino functionalized, cellulose membrane discs containing 9-fluorenylmethyloxycarbonyl-β- alanine (Fmoc-β-Ala) linkers (average loading: 130 nmol/disc – 4 mm diameter). Synthesis was initiated by Fmoc deprotection using 20% piperidine (pip) in dimethylformamide (DMF) followed by washing with DMF and ethanol (EtOH). Peptide chain elongation was achieved using a coupling solution consisting of preactivated amino acids (aas, 0.5 M) with ethyl 2-cyano-2 (hydroxyimino) acetate (oxyma, 1 M) and N,N′-diisopropylcarbodiimide (DIC, 1 M) in DMF (1:1:1, aa:oxyma:DIC). Couplings were carried out x3x30 min, followed by capping (4% acetic anhydride in DMF) and washes with DMF and EtOH. Synthesis was finalized by deprotection with 20% pip in DMF (2×4 μL/disc for 10 min each), followed by washing with DMF and EtOH. Dried discs were transferred to 96 deep-well blocks and treated, while shaking, with sidechain deprotection solution, consisting of 90% trifluoracetic acid (TFA), 2% dichloromethane (DCM), 5% H2O and 3% triisopropylsilane (TIPS) (150 μL/well) for 1.5 h at room temperature (rt). Afterwards, the deprotection solution was removed, and the discs were solubilized overnight at rt, while shaking, using a solvation mixture containing 88.5% TFA, 4% trifluoromethanesulfonic acid (TFMSA), 5% H2O and 2.5% TIPS (250 μL/well). The resulting peptide-cellulose conjugates (PCCs) were precipitated with ice-cold ether (0.7 mL/well) and spun down at 2000×g for 10 min at 4 °C, followed by two additional washes of the formed pellet with ice-cold ether. The resulting pellets were dissolved in DMSO (250 μL/well) to give final stocks. PCC solutions were mixed 2:1 with saline-sodium citrate (SSC) buffer (150 mM NaCl, 15 mM trisodium citrate, pH 7.0) and transferred to a 384-well plate. For transfer of the PCC solutions to white coated CelluSpot blank slides (76×26 mm, Intavis

AG), a SlideSpotter (CEM GmbH) was used. After completion of the printing procedure, slides were left to dry overnight.

**Supplemetary Method 3. Microarray-based probe development**

The microarray contained 241 15aa long peptides representing a full positional scan of the gephyrin protein (GPHN-1 isoform) with a 12aa overlap between the peptides and additional 45 Ser 268/270 phosphorylated peptide versions (Supplementary Table 3). The microarray slides were blocked for 60 min. in 2% (w/v) skimmed milk powder (Carl Roth) phosphate-buffered saline (PBS; 137 mM NaCl, 2.7 mM KCl, 10 mM Na2HPO4, 1.8 mM KH2PO4, pH 7.4). After blocking, the slides were incubated for 30 min. with 1:2500 dilution of mAb7a. mAb7a was detected with a secondary 1:5000 diluted HRP-coupled Anti-mouse antibody (G-21040, Invitrogen). The antibodies were applied in blocking buffer for 30 min., with x3 PBS washes between the antibodies and after the application of the secondary antibody. Chemiluminescent readout was obtained ("High sensitivity" mode (highest resolution; 1536 x 1024), 1s exposure time) after application of 200 µl of SuperSignal™ West Femto Atto Ultimate Sensitivity Substrate (Thermo Fischer Scientific Inc., Waltham, U.S.; Lot: A38554) per slide using a ImageQuant™ LAS 4000 imaging system (GE Healthcare Inc., Chicago, U.S.). Binding intensities were acquired with FIJI using the "Microarray Profile" plugin (OptiNav). The error range and the relative standard deviation were defined by comparing the intensities of each peptide duplicate on the respective array.

### Appendix 1. Gephyrin isoform vector map

**Plasmid map**

pLVX-hSyn-Flag-V5-mScarlett-GPHN-1-IRES-ZsGreen1 (version Eral)

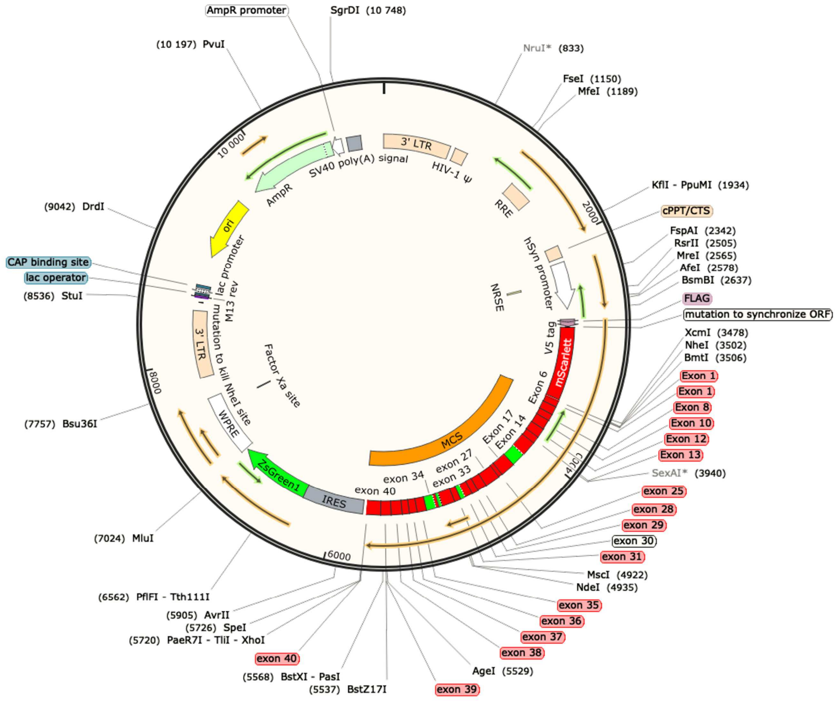

### Appendix 2. Macro and script for Image analysis

| Macro | Mode of operation | Utilization |
|---|---|---|
| I | • renaming the data format | • conversion of the proprietary image format to .tiff<br>• data import |
| II | • averages the image set to one image (noise reduction)<br>• sets greyscale representation of intensities<br>• sets the min and max values for the intensity display | • averaging of image series |
| III | • creates a binary mask and inverses the image<br>• measures the image background and subtracts the background | • subtraction of the image background |
| IV | • opens corresponding channel images iteratively<br>• automatically analyses the images in JACoP plugin and measures the colocalization | • colocalization analysis |
| V | • creates a binary mask corresponding to fluorescent protein location in the cell/object of interest<br>• tracks the binary mask in the target channel and measures the average intensity in ROI | • measurement of intensity in fluorescent protein rich regions in all color channels |

**Macro I:**

```
path = File.openDialog("Select a File");
oldname = File.getName(path);
run("Bio-Formats Macro Extensions");
Ext.setId(path);
Ext.getCurrentFile(file);
Ext.getSeriesCount(seriesCount);
for (s=1; s<=seriesCount; s++) {
	run("Bio-Formats Importer", "open=&path autoscale color_mode=Default view=Hyperstack stack_order=XYCZT series_"+s);
	oldtitle = getTitle();
	newtitle = replace(oldtitle, oldname + " - ", "");
	out_path = getDirectory("image") + newtitle;
	saveAs("tiff", out_path);
	run("Close All");
   }
```

**Macro II:**

```
dir1 = getDirectory("Select directory to average");
list = getFileList(dir1);
SaveDir = getDirectory("Select output directory");
for (g=0; g<list.length; g++) {
	open(dir1 + list[g]);
	originalImageName = getTitle();
	selectWindow(originalImageName);
	run("Z Project...", "projection=[Average Intensity]");
	selectWindow("AVG_" + originalImageName);
	run("Grays");
	//run("Brightness/Contrast...");
	setMinAndMax(150, 2500);
	save(SaveDir + originalImageName);
	close();
	close();
}
```

**Macro III:**

```
dir1 = getDirectory("Select image directory");
list = getFileList(dir1);
SaveDir = getDirectory("Select results directory");
	run("Set Measurements...", "area mean standard min integrated median redirect=Nondecimal=3");
	for (g=0; g<list.length; g++) {
	ch1name = list[g];
	//Identifier = "-Probe";
	print(ch1name);
	open(dir1 + ch1name);
	//rename (g + Identifier);
	//creating a binary mask
	selectWindow(ch1name);
	run("Duplicate...", "title=Dup.tif");
	selectWindow("Dup.tif");
	run("Median...", "radius=5");
	run("Maximum...", "radius=5");
	run("Minimum...", "radius=5");
	setAutoThreshold("Default dark");
	setThreshold(250, 65535);

	setOption("BlackBackground", true);
	run("Convert to Mask");
	run("Make Inverse");
	roiManager("Add");
	RunningNumber = g + 1;
```

```
		roiManager("Save", SaveDir + ch1name + ".roi");
		selectWindow("Dup.tif");
		close();
	//Mask done
	selectWindow(ch1name);
	roiManager("Select", 0);
		run("Measure");
		BG = getResult("Mean");
		print(BG);
		selectWindow(ch1name);
		run("Select None");
		run("Subtract...", "value=BG");
		setMinAndMax(0, 10000);
		save(SaveDir + ch1name);
		close();
		roiManager("reset");
	}
	selectWindow("Results");
	saveAs("Measurements", SaveDir + "Results.tsv");
	selectWindow("ROI Manager");
		run("Close");
	}
```

**Macro IV:**

```
//Coloc measurement 2 folders, JACoP
function parseJACoP() { //Log to table function
	//Get the log window
	logdump = split(getInfo("log"), "\n");
	thrVals = false;
	imgA = -1;
	imgB= -1;
	Pc = -1;
	Oc = -1;
	OcThr = -1;
	k1Thr = -1;
	k1 = -1;
	k2Thr = -1;
	k2 = -1;
	thrA = -1;
	thrB = -1;
	M1 = -1;
	M2 = -1;
	M1Thr = -1;
	M2Thr = -1;
	a = -1;
	b = -1;
	R = -1;
	icq = -1;
	for (i=0; i<logdump.length; i++) {
		if (startsWith(logdump[i], "Image A"))
			imgA = substring(logdump[i], 9, lengthOf(logdump[i]));
		if (startsWith(logdump[i], "Image B"))
			imgB = substring(logdump[i], 9, lengthOf(logdump[i]));
		if (startsWith(logdump[i], "Pearson's Coefficient"))
			Pc = parseFloat(substring(logdump[i+1], 2, lengthOf(logdump[i+1])));

		if (startsWith(logdump[i], "Overlap Coefficient"))
			if (thrVals) {
				OcThr = parseFloat(substring(logdump[i+1], 2, lengthOf(logdump[i+1])));
```

```
				} else {
					Oc = parseFloat(substring(logdump[i+1], 2, lengthOf(logdump[i+1])));
				}
			if (startsWith(logdump[i], "k1=")) {
				if (thrVals) {
					k1Thr = parseFloat(substring(logdump[i], 3, lengthOf(logdump[i])));
				} else {
					k1 = parseFloat(substring(logdump[i], 3, lengthOf(logdump[i])));
				}
			}
			if (startsWith(logdump[i], "k2=")) {
				if (thrVals) {
					k2Thr = parseFloat(substring(logdump[i], 3, lengthOf(logdump[i])));
				} else {
					k2 = parseFloat(substring(logdump[i], 3, lengthOf(logdump[i])));
				}
			}
			if (startsWith(logdump[i], "Using thresholds")) {
				thrA = parseFloat(substring(logdump[i], indexOf(logdump[i], "=")+1, indexOf(logdump[i], "and")-1));
				thrB = parseFloat(substring(logdump[i], lastIndexOf(logdump[i], "=")+1, lastIndexOf(logdump[i], ")")));
				thrVals = true;
			}
			if (startsWith(logdump[i], "Manders' Coefficients (original):")) {
				M1 = parseFloat(substring(logdump[i+1], 3, 8));
				M2 = parseFloat(substring(logdump[i+2], 3, 8));
			}
			if (startsWith(logdump[i], "Manders' Coefficients (using threshold")) {
				M1Thr = parseFloat(substring(logdump[i+1], 3, 8));
				M2Thr = parseFloat(substring(logdump[i+2], 3, 8));
			}
			if (startsWith(logdump[i], "Cytofluorogram's parameters:")) {
				a = parseFloat(substring(logdump[i+1], 3, 8));
				b = parseFloat(substring(logdump[i+2], 3, 8));
				R = parseFloat(substring(logdump[i+3], 25, 30));
			}
			if (startsWith(logdump[i], "ICQ"))
				icq = parseFloat(substring(logdump[i], 5, lengthOf(logdump[i])));
		}
		n=nResults;
		setResult("Pearson's", n, Pc);
		setResult("Overlap Coefficient (no threshold)", n, Oc);
		setResult("k1 (no threshold)", n, k1);
		setResult("k2 (no threshold)", n, k2);
		setResult("M1 (no threshold)", n, M1);
		setResult("M2 (no threshold)", n, M2);
		setResult("ThrA", n, thrA);
		setResult("ThrB", n, thrB);
		setResult("Correlation Coefficient", n, R);
		setResult("Overlap Coefficient", n, OcThr);
		setResult("k1", n, k1Thr);
		setResult("k2", n, k2Thr);
		setResult("M1", n, M1Thr);
		setResult("M2", n, M2Thr);
		setResult("Li's ICQ", n, icq);
		setResult("Cytofluorogram Slope", n, a);
		setResult("Cytofluorogram Intercept", n, b);
}
//Function end
```

```
dir1 = getDirectory("Select Green image directory");
list = getFileList(dir1);
dir2 = getDirectory("Select Far-Red image directory");
SaveDir = getDirectory("Select results directory");
for (g=0; g<list.length; g++) {
        ch1name = list[g];
        RefCh = "R";
        TargetCh = "FR";
        ch2name = replace(ch1name, RefCh, TargetCh)
        open(dir1 + ch1name);
        open(dir2 + ch2name);
        run("JACoP ", "imga=["+ch1name+"] imgb=["+ch2name+"] thra=400 thrb=160 pearson mm");
        parseJACoP();
        // Clear log window
        print("\\Clear");
        close(ch1name);
        close(ch2name);
}
saveAs("Measurements", SaveDir + "ColocData.tsv");
```

**Macro V:**

```
dir1 = getDirectory("Select reference image directory");
list = getFileList(dir1);
dir2 = getDirectory("Select target image directory");
SaveDir = getDirectory("Select output directory");
for (g=0; g<list.length; g++) {
        RunningNumber = g + 1;
        ch1name = list[g];
        RefCh = "R";
        TargetCh = "FR";
        ch2name = replace(ch1name, RefCh, TargetCh);
        print(ch1name);
        open(dir1 + ch1name);
//creating a Mask from a reference channel ROI [R]
        selectWindow(ch1name);
        run("Duplicate...", "title=Dup.tif");
        selectWindow("Dup.tif");
        run("Median...", "radius=5");
        run("Maximum...", "radius=5");
        run("Minimum...", "radius=5");
        setAutoThreshold("Default dark");
        setThreshold(600, 65535);
        setOption("BlackBackground", true);
        run("Convert to Mask");
        run("Create Selection");
        roiManager("Add");
        roiManager("Save", SaveDir + RunningNumber + ".roi");
        selectWindow("Dup.tif");
        close();
//Mask done
//creating 2nd Mask
        selectWindow(ch1name);
        run("Duplicate...", "title=Dup.tif");
        selectWindow("Dup.tif");
        setAutoThreshold("Default dark");
        setThreshold(4000, 65535);
        setOption("BlackBackground", true);
        run("Convert to Mask");
        run("Create Selection");
        roiManager("Add");
```

```
        roiManager("Select", 1);
        roiManager("Save", SaveDir + RunningNumber + "-punctae" + ".roi");
        selectWindow("Dup.tif");
        close();
//Mask done
//3rd mask for mScarlet-gephyrin controls
        selectWindow(ch1name);
        run("Duplicate...", "title=Dup.tif");
        selectWindow("Dup.tif");
        run("Median...", "radius=5");
        run("Maximum...", "radius=5");
        run("Minimum...", "radius=5");
        setAutoThreshold("Default dark");
        setThreshold(1000, 65535);
        setOption("BlackBackground", true);
        run("Convert to Mask");
        run("Create Selection");
        roiManager("Add");
        roiManager("Select", 2);
        roiManager("Save", SaveDir + RunningNumber + "-foreGFP" + ".roi");
        selectWindow("Dup.tif");
        close();
        //Mask done
        selectWindow(ch1name);
        roiManager("Select", 0);
        run("Measure");
        roiManager("Select", 1);
        run("Measure");
        roiManager("Select", newArray(0,1));
        roiManager("XOR");
        run("Measure");
        roiManager("Select", 2);
        run("Measure");
        close();
        open(dir2 + ch2name);
        roiManager("Select", 0);
        run("Measure");
        roiManager("Select", 1);
        run("Measure");
        roiManager("Select", newArray(0,1));
        roiManager("XOR");
        run("Measure");
        roiManager("Select", 2);
        run("Measure");
        close();
        roiManager("reset");
}
selectWindow("Results");
saveAs("Measurements", SaveDir + "Results.tsv");
run("Close");
selectWindow("ROI Manager");
run("Close");
```

### Appendix 3. Icy 2.0.3.0 protocol for single synapse segmentation and intensity recording

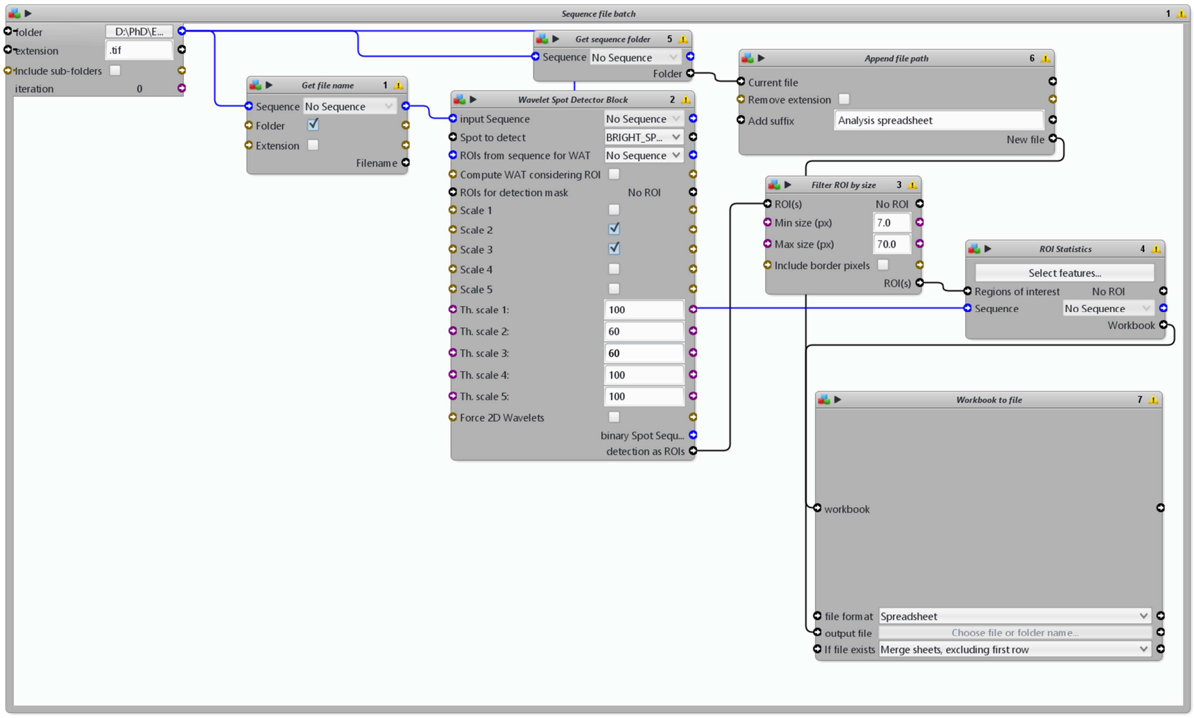

### Appendix 4. Spectral profiles of Sylites

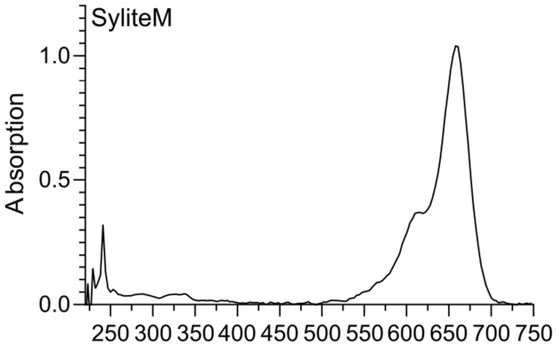

**Excitation**

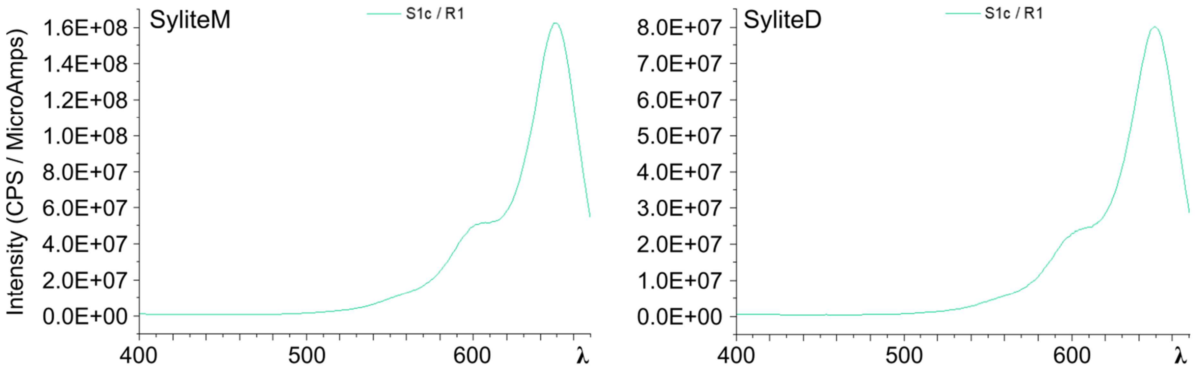

**Emission**

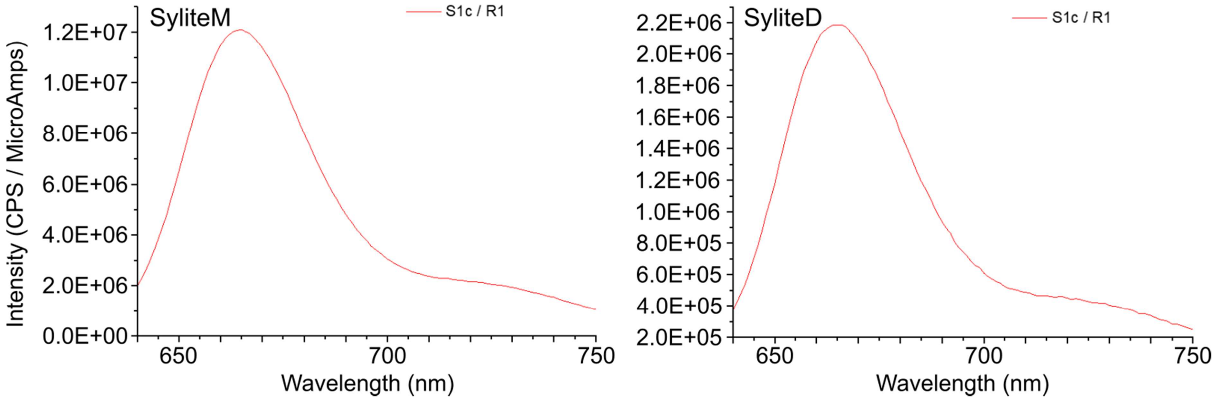

**Appendix 5. Chromatographic and mass spectrometric probe validation**

**SyliteM (VK15)**

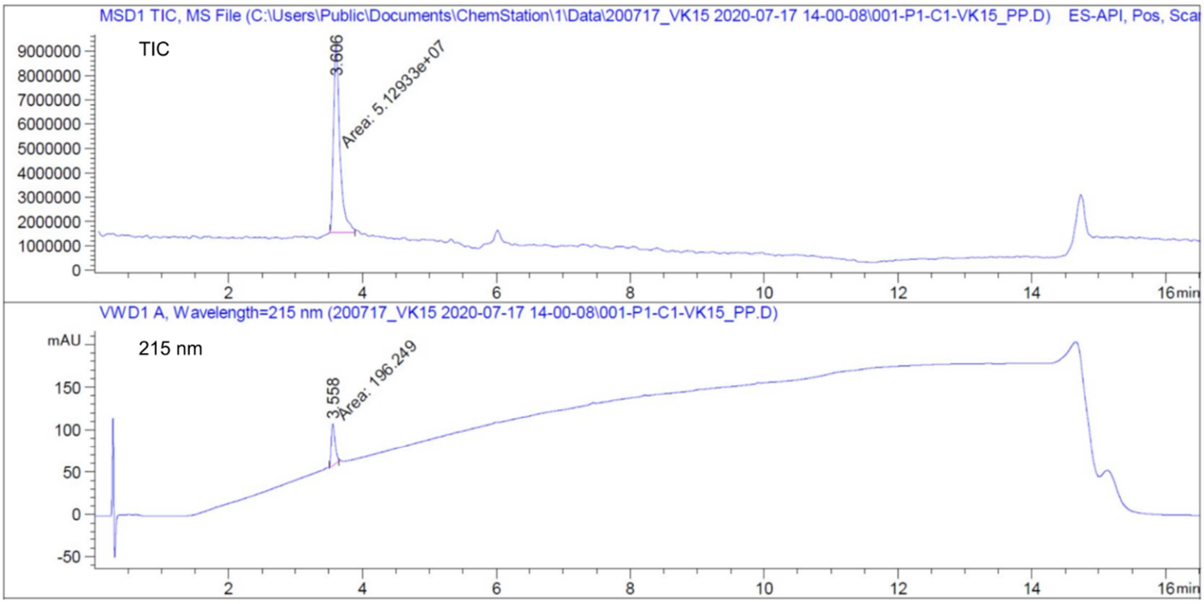

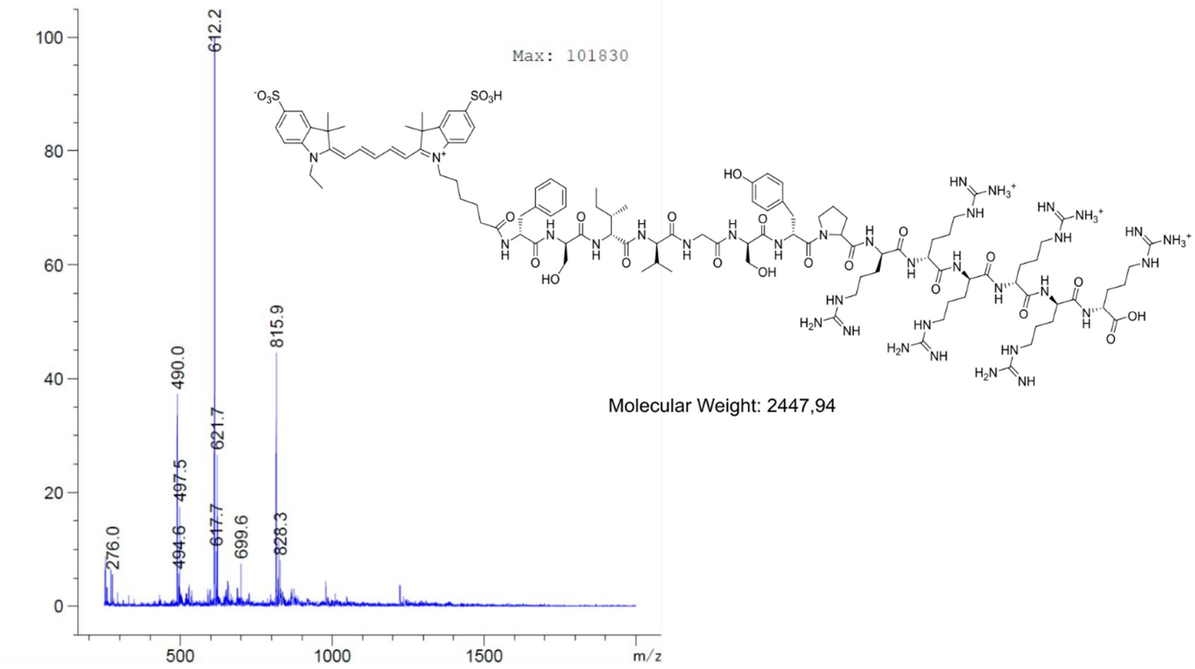

**SyliteD (NN1D)**

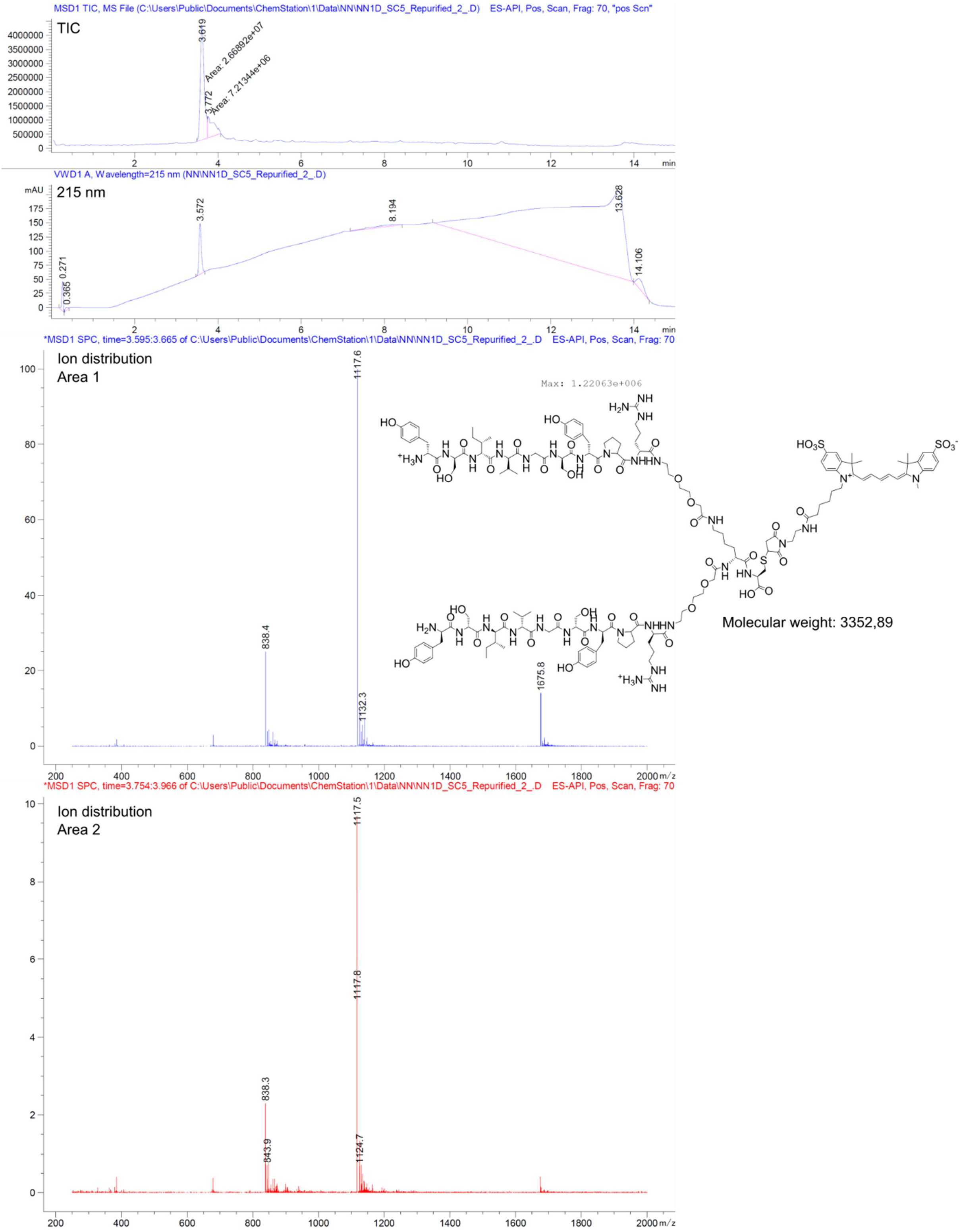

**VK16**

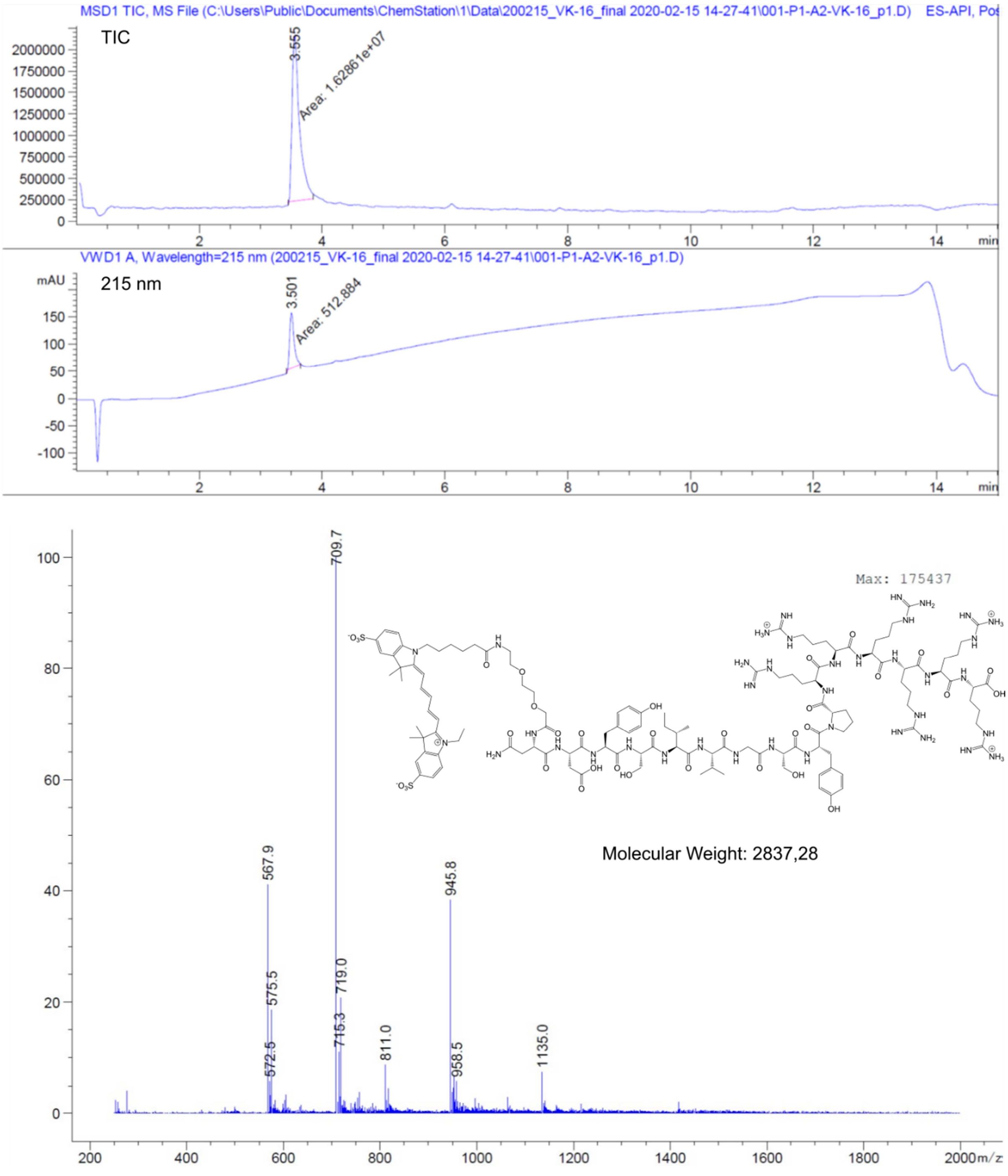

MSD1 TIC, MS File (C:\Users\Public\Documents\ChemStation\1\Data\200215_VK-16_final 2020-02-15 14-27-41\001-P1-A2-VK-16_p1.D) ES-API, Pos
TIC
3.555
Area: 1.62861e+07
VWD1 A, Wavelength=215 nm (200215_VK-16_final 2020-02-15 14-27-41\001-P1-A2-VK-16_p1.D)
215 nm
mAU
3.501
Area: 512.884
min
709.7
567.9
575.5
572.5
715.3
719.0
811.0
945.8
958.5
1135.0
Max: 175437
Molecular Weight: 2837,28
m/z

**VK17**

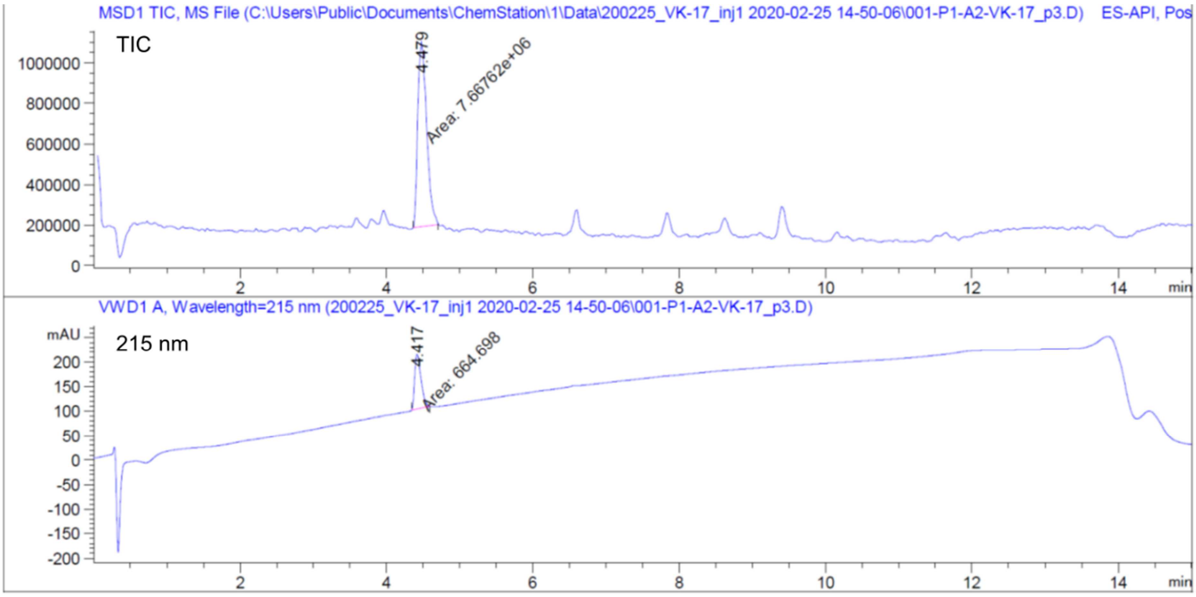
MSD1 TIC, MS File (C:\Users\Public\Documents\ChemStation\1\Data\200225_VK-17_inj1 2020-02-25 14-50-06\001-P1-A2-VK-17_p3.D) ES-API, Pos
TIC
4.479
Area: 7.66762e+06
VWD1 A, Wavelength=215 nm (200225_VK-17_inj1 2020-02-25 14-50-06\001-P1-A2-VK-17_p3.D)
mAU
215 nm
4.417
Area: 664.698
min

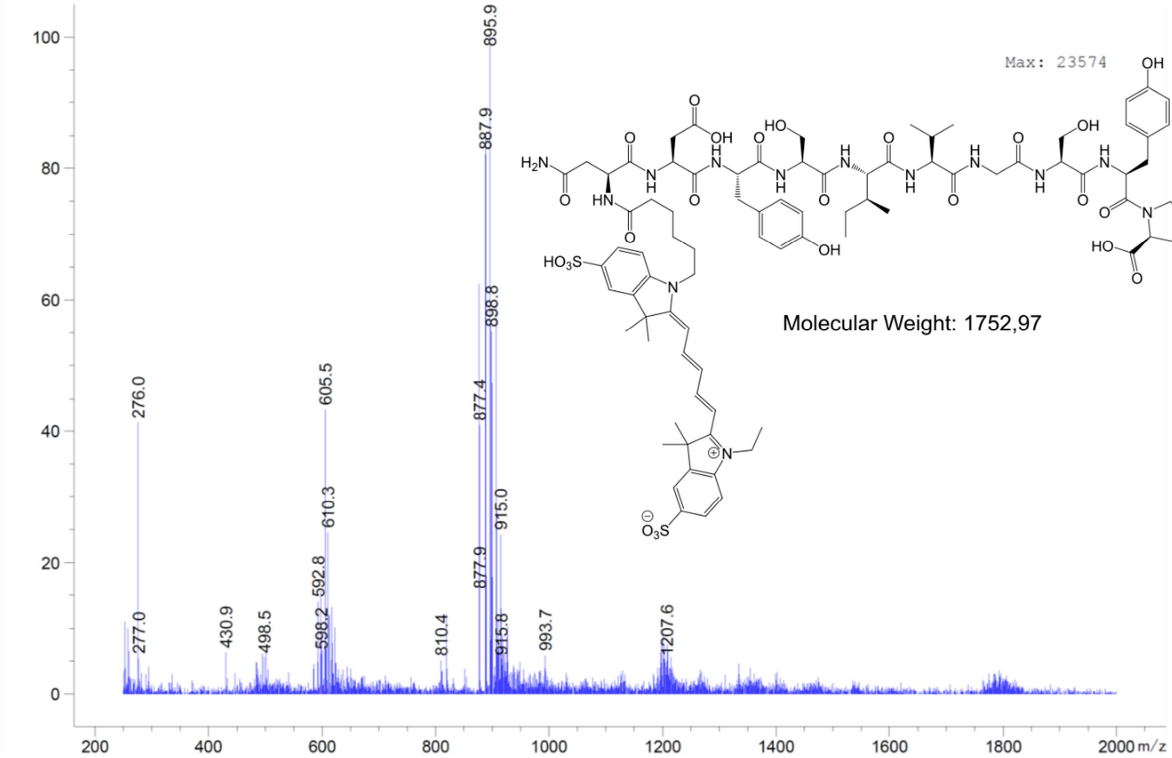
Max: 23574
Molecular Weight: 1752,97
895.9
887.9
898.8
877.4
915.0
877.9
276.0
605.5
610.3
592.8
598.2
277.0
430.9
498.5
810.4
915.8
993.7
1207.6
m/z

**VK18**

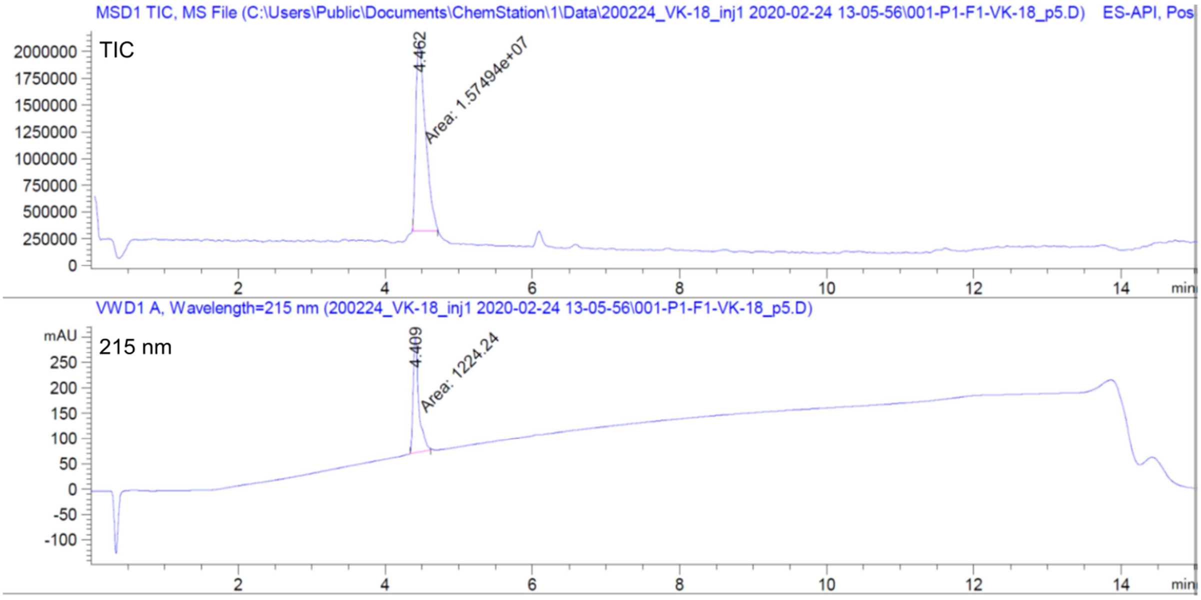

MSD1 TIC, MS File (C:\Users\Public\Documents\ChemStation\1\Data\200224_VK-18_inj1 2020-02-24 13-05-56\001-P1-F1-VK-18_p5.D) ES-API, Pos
TIC
4.462
Area: 1.57494e+07
VWD1 A, Wavelength=215 nm (200224_VK-18_inj1 2020-02-24 13-05-56\001-P1-F1-VK-18_p5.D)
mAU
215 nm
4.409
Area: 1224.24
min

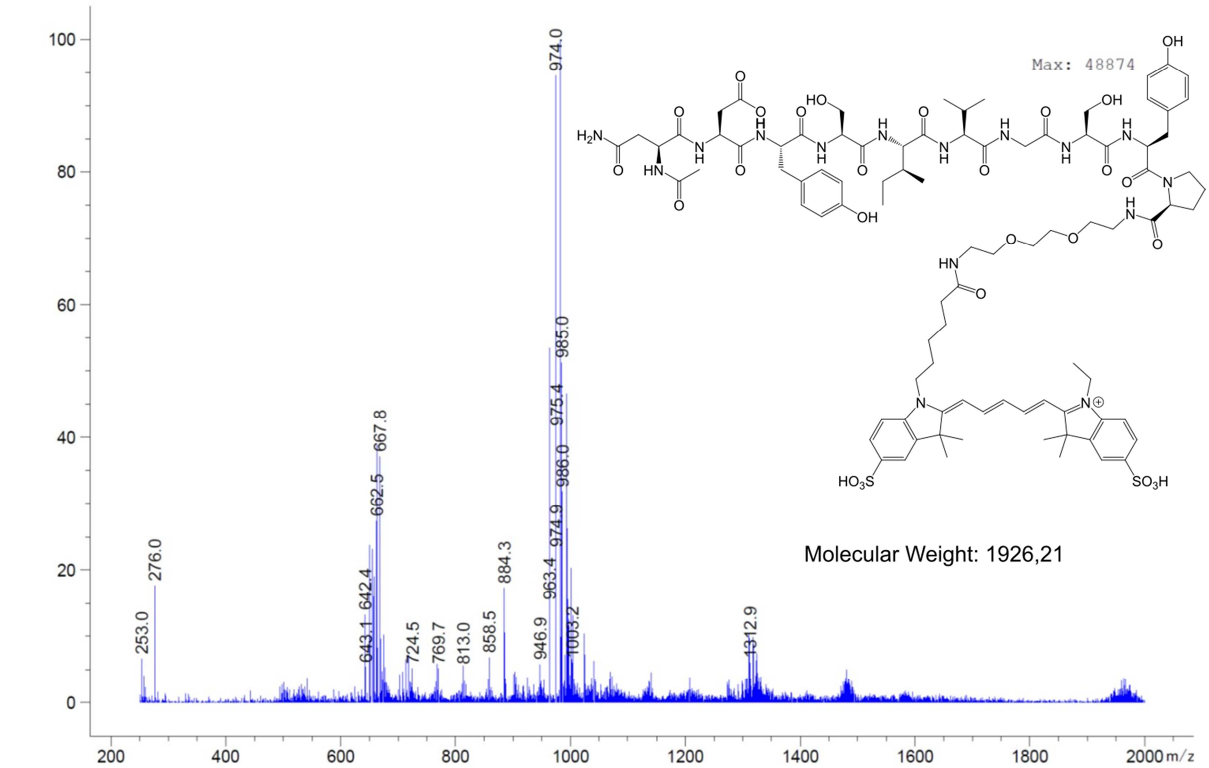

Max: 48874
974.0
985.0
975.4
667.8
986.0
662.5
974.9
884.3
276.0
963.4
642.4
1312.9
253.0
1003.2
643.1
724.5
769.7
813.0
858.5
946.9
m/z
Molecular Weight: 1926,21

**VK20**

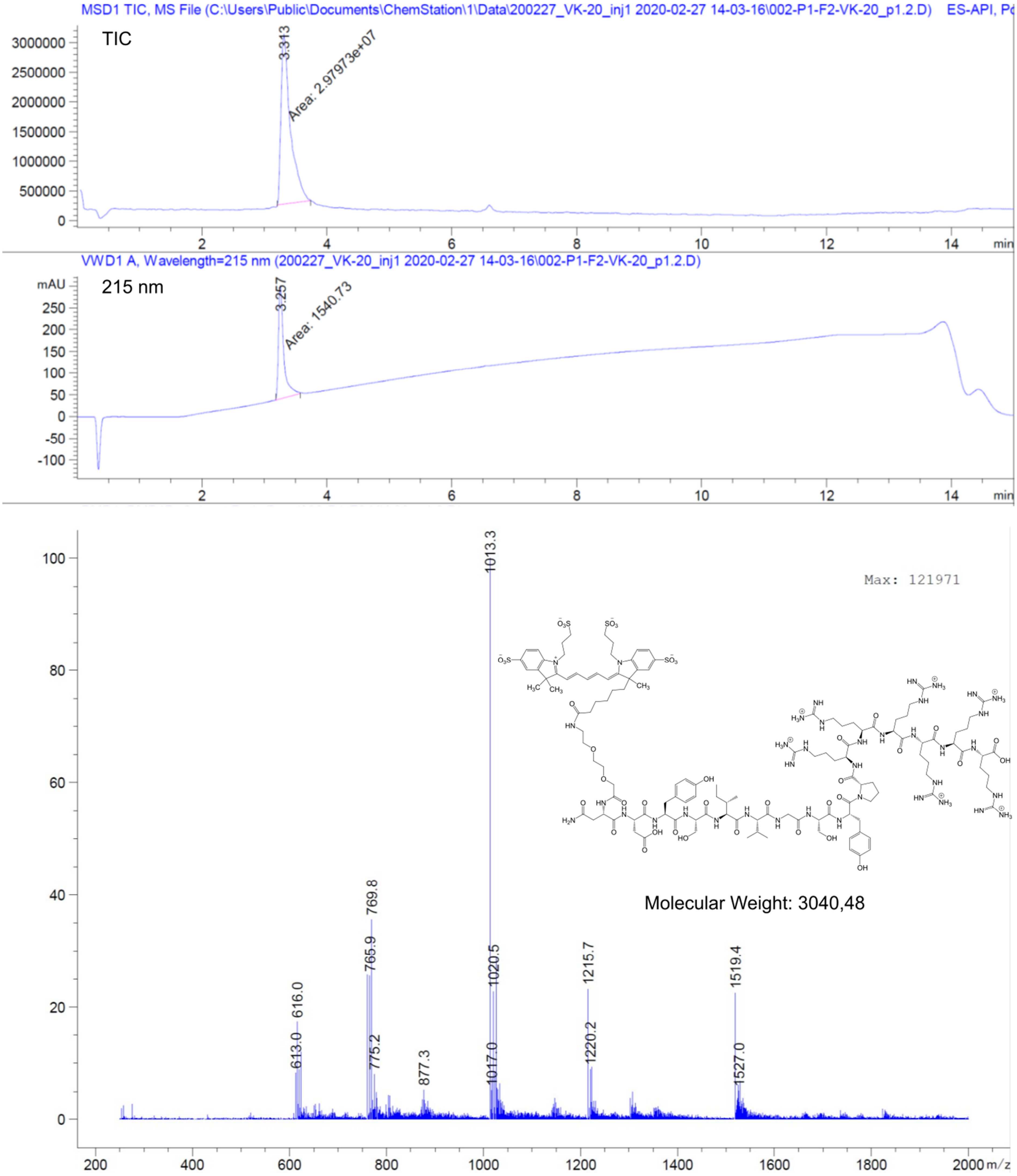

MSD1 TIC, MS File (C:\Users\Public\Documents\ChemStation\1\Data\200227_VK-20_inj1 2020-02-27 14-03-16\002-P1-F2-VK-20_p1.2.D) ES-API, Po
TIC
3.313
Area: 2.97973e+07
VWD1 A, Wavelength=215 nm (200227_VK-20_inj1 2020-02-27 14-03-16\002-P1-F2-VK-20_p1.2.D)
215 nm
mAU
3.257
Area: 1540.73
min
Max: 121971
1013.3
769.8
765.9
1020.5
1215.7
1519.4
616.0
613.0
775.2
877.3
1017.0
1220.2
1527.0
Molecular Weight: 3040,48
m/z

**VK21**

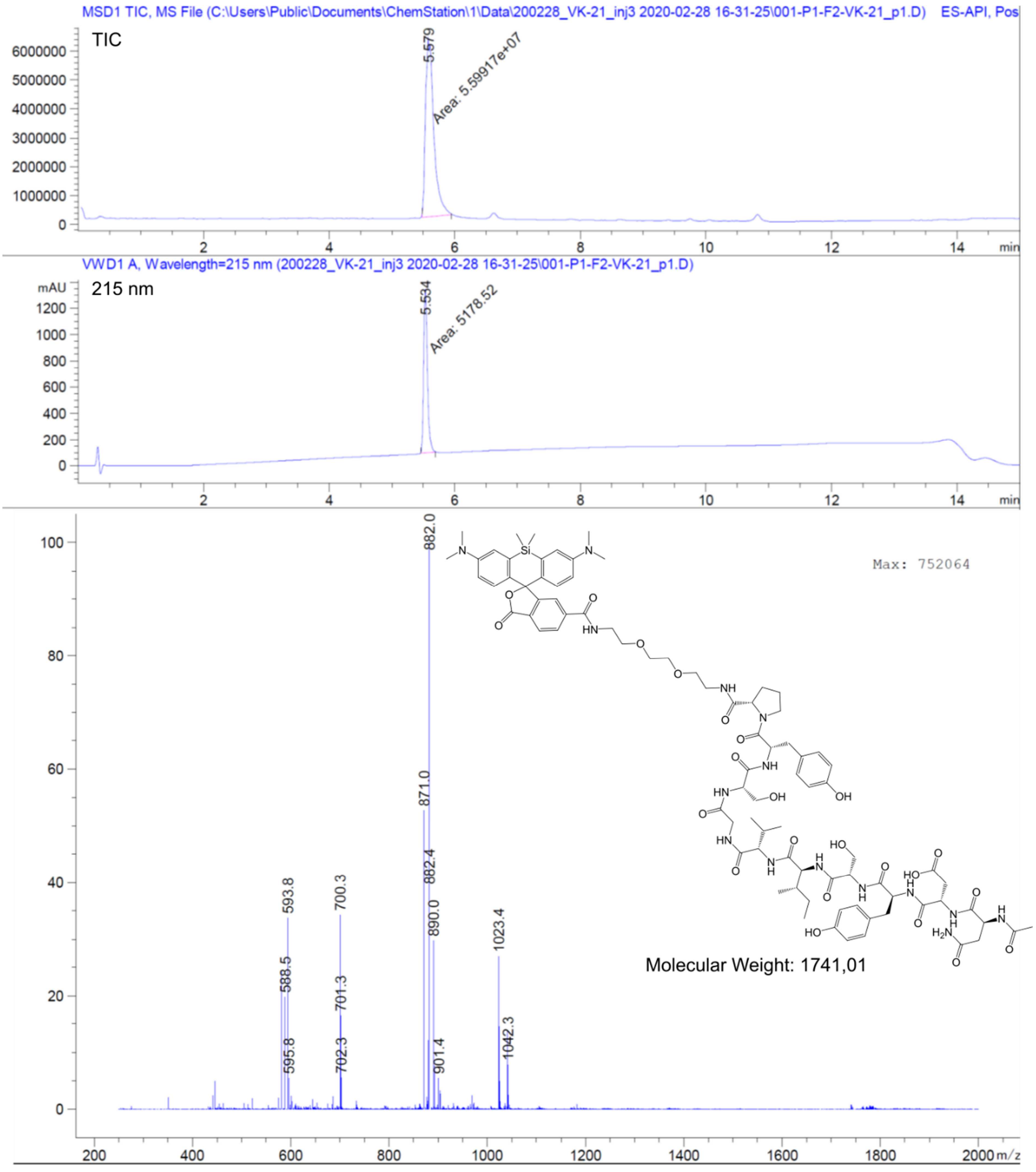
MSD1 TIC, MS File (C:\Users\Public\Documents\ChemStation\1\Data\200228_VK-21_inj3 2020-02-28 16-31-25\001-P1-F2-VK-21_p1.D) ES-API, Pos
TIC
5.579
Area: 5.59917e+07
VWD1 A, Wavelength=215 nm (200228_VK-21_inj3 2020-02-28 16-31-25\001-P1-F2-VK-21_p1.D)
215 nm
mAU
5.534
Area: 5178.52
min
Max: 752064
882.0
871.0
882.4
890.0
593.8
588.5
595.8
700.3
701.3
702.3
901.4
1023.4
1042.3
m/z
Molecular Weight: 1741,01

**VK22**

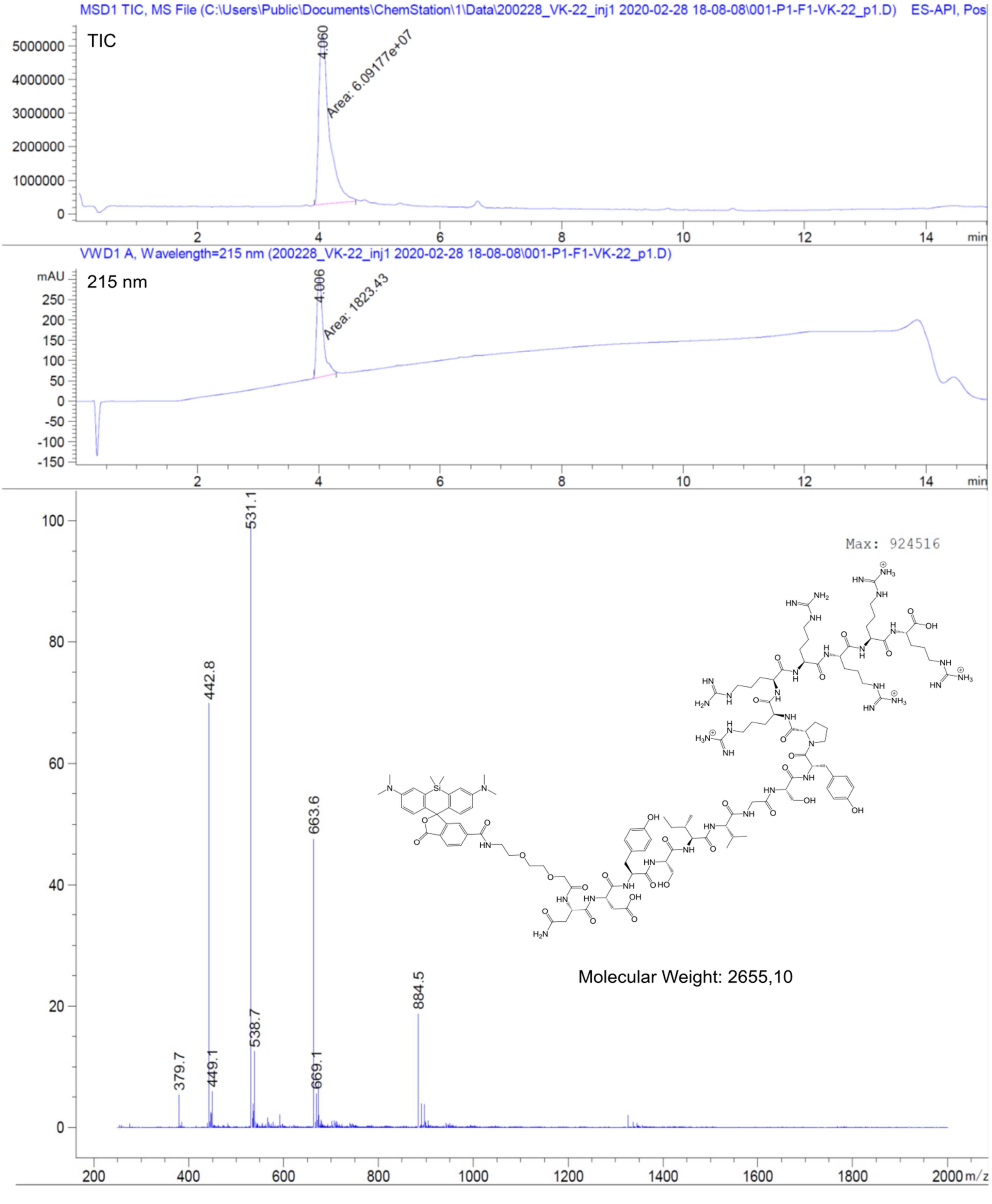
MSD1 TIC, MS File (C:\Users\Public\Documents\ChemStation\1\Data\200228_VK-22_inj1 2020-02-28 18-08-08\001-P1-F1-VK-22_p1.D) ES-API, Pos
TIC
4.060
Area: 6.09177e+07
min
VWD1 A, Wavelength=215 nm (200228_VK-22_inj1 2020-02-28 18-08-08\001-P1-F1-VK-22_p1.D)
mAU
215 nm
4.006
Area: 1823.43
min
Max: 924516
531.1
442.8
663.6
884.5
538.7
379.7
449.1
669.1
m/z
Molecular Weight: 2655,10

**DVK1**

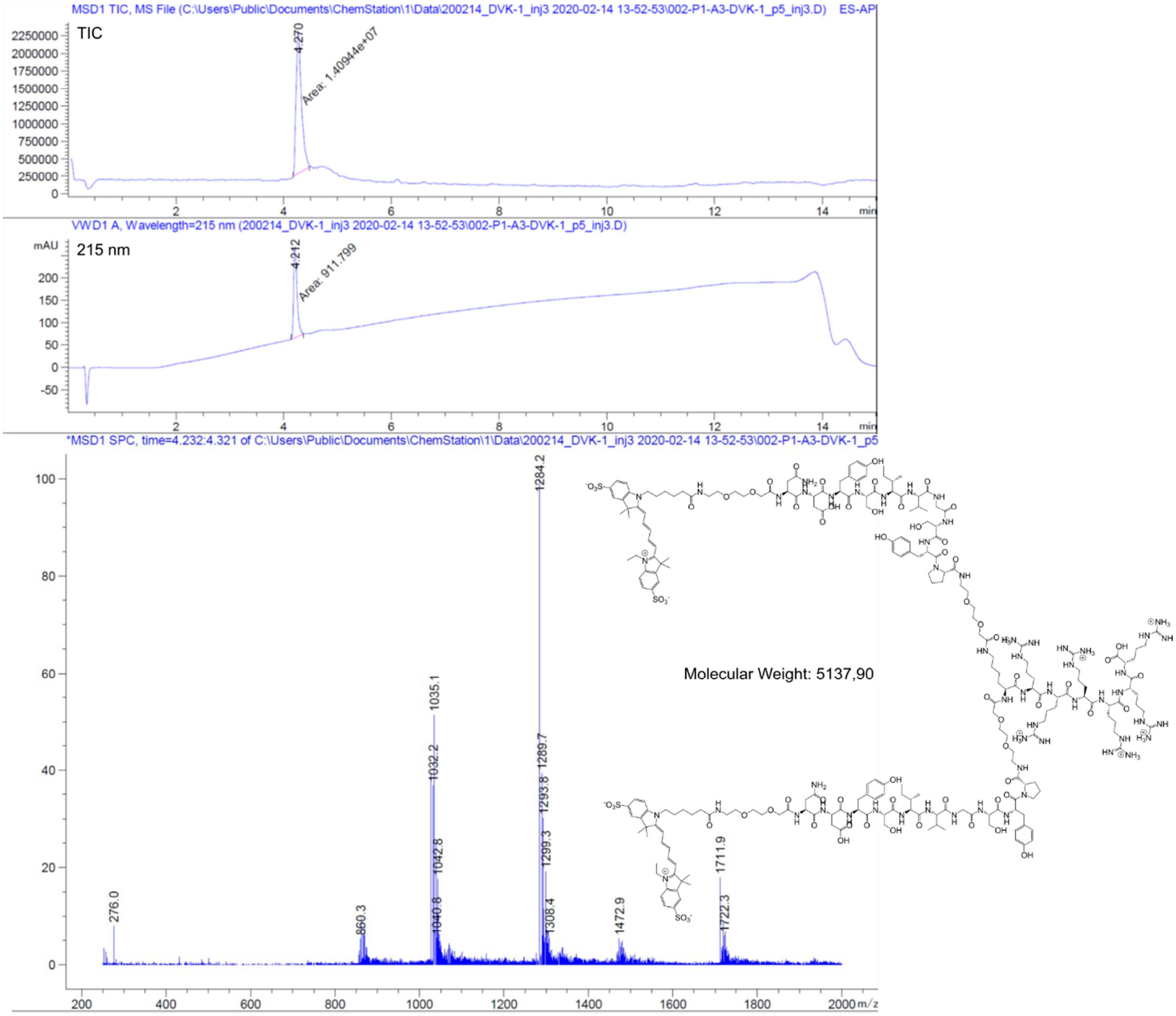
MSD1 TIC, MS File (C:\Users\Public\Documents\ChemStation\1\Data\200214_DVK-1_inj3 2020-02-14 13-52-53\002-P1-A3-DVK-1_p5_inj3.D) ES-API
TIC
4.270
Area: 1.40944e+07
min
VWD1 A, Wavelength=215 nm (200214_DVK-1_inj3 2020-02-14 13-52-53\002-P1-A3-DVK-1_p5_inj3.D)
mAU
215 nm
4.212
Area: 911.799
min
*MSD1 SPC, time=4.232:4.321 of C:\Users\Public\Documents\ChemStation\1\Data\200214_DVK-1_inj3 2020-02-14 13-52-53\002-P1-A3-DVK-1_p5
1284.2
1035.1
1032.2
1289.7
1293.8
1042.8
1299.3
1040.8
1308.4
860.3
276.0
1472.9
1711.9
1722.3
m/z
Molecular Weight: 5137,90

**DVK2**

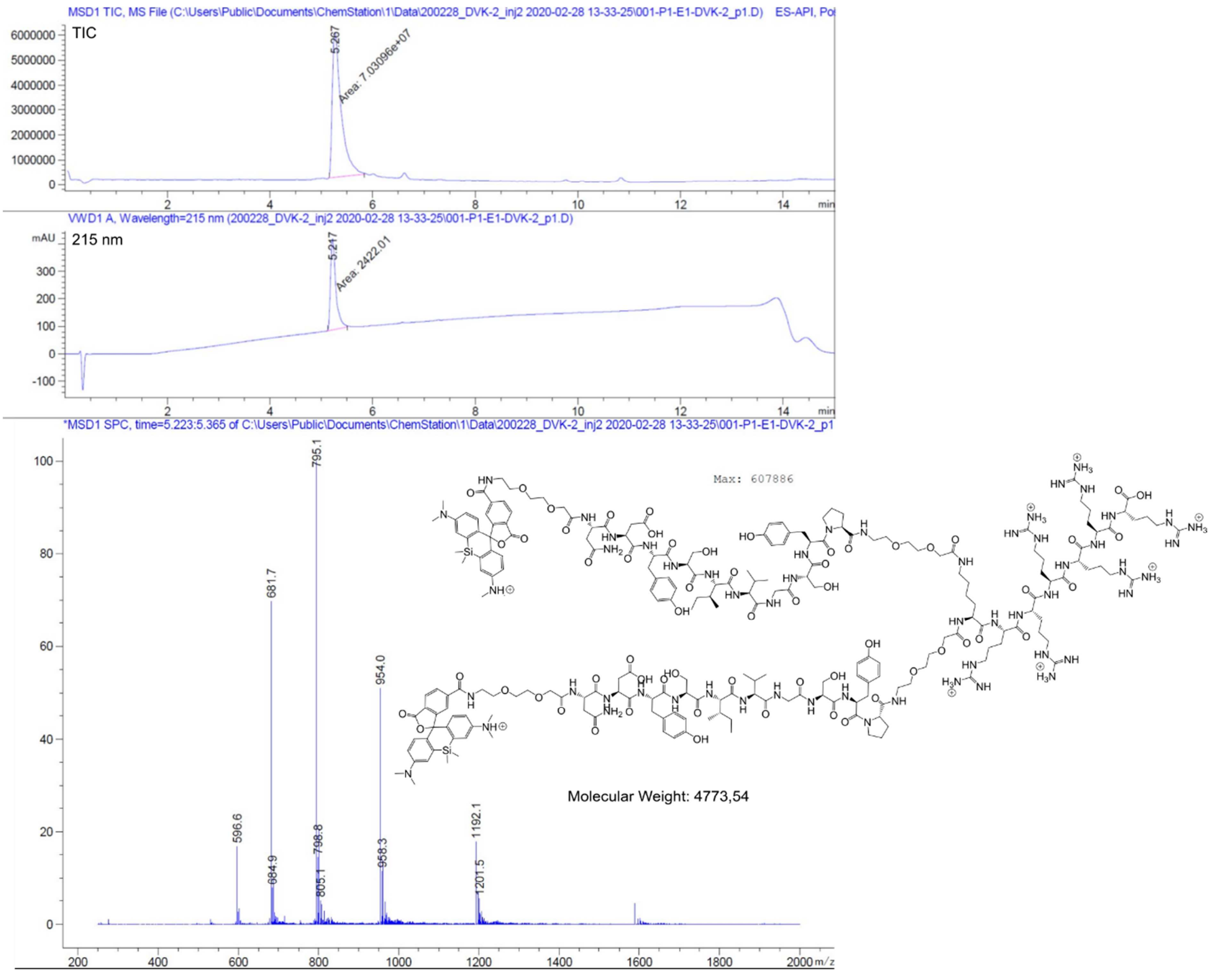

MSD1 TIC, MS File (C:\Users\Public\Documents\ChemStation\1\Data\200228_DVK-2_inj2 2020-02-28 13-33-25\001-P1-E1-DVK-2_p1.D) ES-API, Po
TIC
5.267
Area: 7.03096e+07
VWD1 A, Wavelength=215 nm (200228_DVK-2_inj2 2020-02-28 13-33-25\001-P1-E1-DVK-2_p1.D)
mAU
215 nm
5.217
Area: 2422.01
min
*MSD1 SPC, time=5.223:5.365 of C:\Users\Public\Documents\ChemStation\1\Data\200228_DVK-2_inj2 2020-02-28 13-33-25\001-P1-E1-DVK-2_p1
Max: 607886
795.1
681.7
954.0
596.6
684.9
798.8
805.1
958.3
1192.1
1201.5
m/z
Molecular Weight: 4773,54
